# Cell type-specific gene regulatory network inference from single cell transcriptomics with ctOTVelo

**DOI:** 10.64898/2026.03.11.711174

**Authors:** Seowon Chang, Wenjun Zhao, Ying Ma, Bjorn Sandstede, Ritambhara Singh

## Abstract

Inferring gene regulatory networks (GRNs) from gene expression is a crucial task for understanding functional relationships. Gene expression data (transcriptomics) provide a snapshot of gene activity, encoding information about gene regulatory relationships. However, gene regulation is a dynamic process, modulating across time and with different cell types. Temporal GRN inference methods aim to capture these dynamics by utilizing time-stamped transcriptomics, gene expression data of similar samples captured across discrete timepoints, or pseudotime transcriptomics, computationally ordering cells based on an inferred trajectory. These methods can estimate constant or temporal gene regulatory relationships, but may not capture finer, cell type specific relationships. We propose ctOTVelo, an extension to our previous work to account for cell type specificity during GRN inference. ctOTVelo incorporates cell type labels or proportions when inferring the GRN from single cell transcriptomics data. Our methods achieve state-of-the-art performance in GRN prediction in time-stamped and pseudotime-stamped transcriptomics. Furthermore, ctOTVelo is able to generate cell type specific GRNs, allowing cell type resolution analysis of gene regulatory relationships.

## 1 Introduction

Gene regulatory networks (GRNs) describe intricate biological mechanisms. How genes regulate each other and the strength of their regulatory effect are critical to describing complex processes such as cell fate, environmental response, or developmental processes. Recent approaches have utilized single cell RNA-sequencing (scRNA-seq)to infer GRNs. Single-cell technologies allow observations at the single cell resolution, increasing the number of samples measured and revealing cellular heterogeneity. Furthermore, advances in scRNA-seq allow the profiling of cells across discrete time points. Alternatively, pseudotimes can be computed for scRNA-seq, which orders cells along predicted trajectories. Given single cell resolution that captures cell-level gene activity and temporal resolution, there is great potential to extract gene regulatory relationships.

Numerous approaches have been proposed to learn GRNs from scRNA-seq datasets. Tree based models, such as GENIE3/GRNBoost [2]and its improved version GENIE3+/GRNBoost2 [12], predict regulatory relationships by predicting gene expression. Regression based models are also popular - for example, SINCERITIES [15] utilizes temporal changes and Granger causality to predict gene regulation. Other methods account for cell lineages when considering regulatory relationships; scMTNI [22] uses a cell lineage tree to infer different GRNs per lineage timepoint. Finally, other methods account for cell-level mismatches between time points. For example, OTVelo [23] utilizes optimal transport to model the cell-level probability mappings between timepoints, estimate gene velocities across time, and compare gene velocities to infer gene regulatory relationships. These approaches demonstrate the strength of inferring gene regulatory relationships from single cell datasets.

While temporal effects on gene regulation have been studied from various angles, one relatively understudied aspect is cell type specificity in gene regulation. Different cell types have heterogeneous biological functions, which are driven by different pathways and correspondingly different gene regulatory patterns. Therefore, different cell types will have different underlying GRNs. For example, studies have shown that cell fate is driven by heterogeneous cell cycles, where pluripotent stem cells show differences in cell cycle regualtion [9]. While one method called scMTNI [22] approaches this from the view of cell lineage-specific GRNs, it cannot capture cell type-specific regulatory patterns as cell lineages consist of a mixture of cell types. A natural direction in GRN inference is then to discern these cell type-specific regulatory relationships. Given the prevalance of single cell data and cell type information associated with those datasets, it is also natural to focus on a model that relies on solely transcriptomics modalities, rather than paired multiomics datasets.

To address this aim, we propose ctOTVelo, a simple framework for identifying cell type-specific GRNs. This work is motivated by previous approaches, particularly OTVelo, which capture cell level and temporal changes but do not take into account cell type information. OTVelo utilizes cell-level probability mappings to estimate rates of change in genes between timepoints, or pseudo-gene velocities. These gene velocities can then be used to calculate time-lagged correlations between genes that suggest pairwise gene regulatory relationships. We propose a modified time-lagged correlation which incorporates cell type information to weight time-lagged relationships between rates of change of gene expression. Because time-lagged correlation utilizes cell type proportions and not specific cell type information, this framework is flexible for all cell types. In summary, we make the following contributions:

- Incorporation of cell type proportions to estimate cell type-drive gene velocities for cell-type specific GRN inference.
- Real dataset analysis of human hematopoietic differentiation and spatiotemporal mouse organogenesis shows state of the art GRN prediction performance.
- Downstream analysis of inferred cell type specific GRNs reveal cell type specific regulatory relationships.

## 2 Related Work

Several algorithms have been proposed to infer GRNs from transcriptomics datasets. For example, SCENIC [2] is a workflow that utilize GENIE3/GENIE3+ and GRNBoost/GRNBoost2 [12], which are tree-based algorithms for computing GRNs. GENIE3 uses random forest models to predict the expression of each gene. The weights for transcription factors derived from these trained models then determine the regulatory strength of each gene, where higher weights denote greater relevance for the transcription factor in predicting a target gene. GRNBoost utilizes the same concept, except with gradient boosted stumps. While GENIE3 and GRNBoost are effective at capturing gene expression level relationships, they do not take temporal or pseudotime information into account, which can be relevant since certain regulatory relationships are time-dependent. SINCERITIES [15] addresses this issue by utilizing a ridge regression and Granger causality to determine gene-gene relationships from temporal changes in gene expression. Benchmarking studies showed that these popular frameworks are generally better at performing on synthetic datasets rather than real experimental data [16]. This motivated methods to consider additional aspects of single cell data to more accurately capture gene regulation. For example, scMTNI [22] utilizes a cell lineage tree to learn GRNs for a given cell lineage time point rather than a summary GRN. OTVelo [23] is an optimal transport based method that considers the cell mismatch problem between time points, since scRNA-seq is a destructive method and does not have one-to-one mappings of cells between time points. However, all of these methods do not incorporate cell type specific dynamics and thus can miss nuances with cell type specific regulatory relationships. Learning global or cell lineage GRNs miss cell type specific relationships and cannot distinguish if gene regulatory relationships are shared or cell type-specific, which loses resolution in downstream analyses. Our work aims to address this gap by inferring cell type specific GRNs without losing overall predictive power.

## 3 Methods

In this section, we introduce ctOTVelo, a method for inferring cell type specific GRNs from RNA-seq data (Figure 1A). As an overview, ctOTVelo first learns a probabilistic mapping between cells of adjacent timepoints through optimal transport (Figure 1B). ctOTVelo then utilizes a cell type-aware time-lagged correlation to infer cell type specific correlations between genes (Figure 1C). The aggregated correlation then can be interpreted as a cell type specific GRN (Figure 1D).

**Figure 1:**
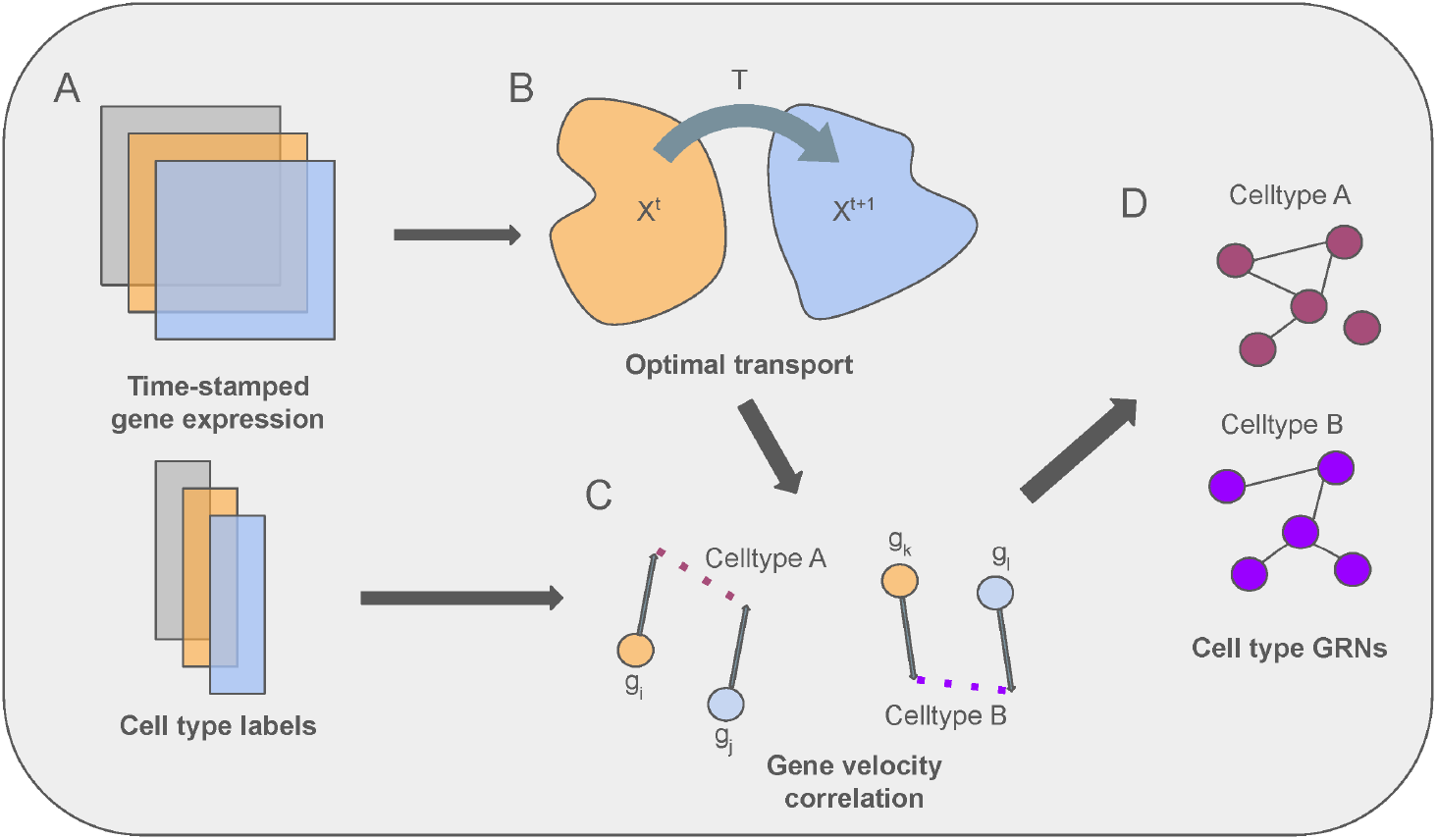
Overview of ctOTVelo. A. Time- or pseudotime-stamped single cell RNA sequencing gene expression data and cell type labels serve as input. B. ctOTVelo learns cell probabilistic mappings between timepoints. C. Using the learned mappings, ctOTVelo computes gene velocities and utilizes time-lagged correlation with cell types. D. From cell type time-lagged correlations between each pair of genes, ctOTVelo can infer cell type-specific GRNs.

### 3.1 Problem Setting

We consider a time-stamped or pseudo-time-stamped gene expression dataset 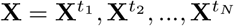, where *N* is the number of discrete timepoints. Each timepoint’s expression data is given as 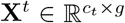, where *c*_*t*_ is the number of cells measured at time *t* and *g* is the number of genes measured across all timepoints. For each cell *c* at timepoint *t*, we also have cell type labels *η*^*c,t*^ ∈ ℝ^*H*^ that can be represented as a vector where 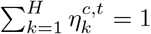, where *H* is the number of cell types. The aim is to infer cell type specific gene regulatory networks 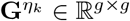, where *G*_*ij*_ indicates a regulatory relationship between genes *i* to *j*.

### 3.2 Inferring cell type specific GRNs with ctOTVelo

Here we describe the framework of ctOTVelo, which utilizes cell type specific information to emphasize cell type specific relationships detected by OTVelo.

#### 3.2.1 Learning temporal cell trajectory with optimal transport

Given two count matrices at consecutive time points *t* and 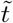, optimal transport learns the transition matrix that measures the probability that a cell from time 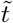 is descended from cells observed in time *t*. We follow the original approach from OTVelo to learn the transition matrix, which uses entropic fused Gromov–Wasserstein optimal transport on unlabeled gene expression data. For continuity, we will utilize OTVelo’s notation. The transition matrix is found by optimizing

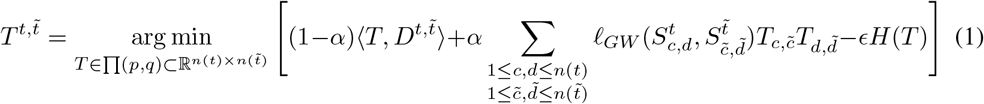

where

- 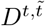 measures the pairwise distances between cells across time points
- 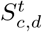 captures the differences between cells *c* and *d* at time *t*
- *ℓ*_*GW*_ is the loss function accounting for misfit distances between the two spaces
- *p* is the distribution of the source space
- *q* is the distribution of the target space
- Π (*p, q*) is the family of transition matrices with the marginals *p* and *q*
- *α* controls the weighting between the Wasserstein and Gromov-Wasserstein terms

The optimal transport optimization problem is solved using projected gradient descent.

For a given cell *c* at time *t*, we can predict its descendant at time 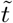 using a barycentric projection using its gene expression *x*^*t,c*^:

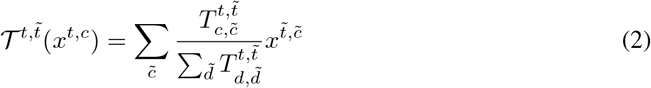

Full notation details are provided in Appendix A.

#### 3.2.2 Identifying gene velocities from temporal cell maps

We can utilize the learned transition matrices 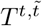 and corresponding barycentric projection map 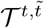 as a measure of cell dynamics moving between time points. Following the original approach from OTVelo, we can then define gene velocity, or the rate of change of gene expression. Following the same notation as OTVelo for clarity, we can define the following velocities for gene expression *x*:

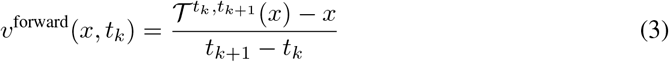

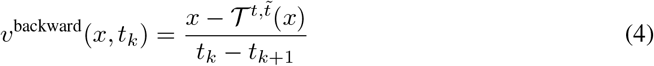

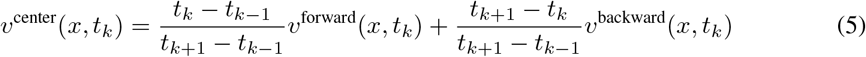

For the first time point, we utilize the forward velocity; for the last time point, we utilize the backward velocity; for all other intermediate points, we utilize center velocity.

#### 3.2.3 Identifying cell type specific gene regulatory relationships with time lagged correlation

To identify gene regulatory relationships, we define time lagged correlation using the computed gene velocities between genes *g*_1_ and *g*_2_. Formally,

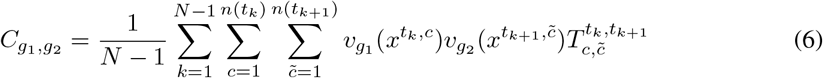

where 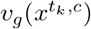 represents the velocity of gene *g* given cell *c* at time *t*. The velocities are normalized to have unit standard deviation across all time points to accounts for varying gene expression levels. For given cell type proportion *L*(*c, η*_*k*_) of cell *c* for cell type *η*_*k*_, we can then define cell type specific regulatory relationships as

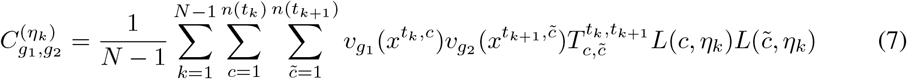

Note that not only are velocities weighted by the probability of transitioning between cells *c* and 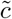, they are also weighted by the cell type proportions. This emphasizes gene interactions for a given cell type *η*_*k*_, thus generating cell-type specific regulatory relationships.

The correlation matrices *C* and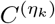 can then be treated as inferred global and cell type specific GRNs respectively.

## 4 Results

In this section, we present our experiments and results evaluating ctOTVelo on two transcriptomics datasets.

### 4.1 Data

We evaluate on a human hematopoiesis differentiation single cell RNA-seq (HHD) [5] and a spatiotemporal mouse organogenesis datasets (SMO) [6]. We choose these datasets for several reasons. First, the HHD dataset is not time-stamped while SMO is temporally resolved, allowing us to test ctOTVelo on both pseudotime- and real time-stamped single cell data. Second, there exists ground truth GRNs through ChIP-seq and perturbation experiments for hematopoiesis and mouse embryos. Finally, these datasets have distinct cell types/states in which downstream cell type specific GRNs should contain differentiating signal.

**Table 1:** Datasets analyzed in the paper.

| Name | Modality | Number of Cells | Number of Genes | Number of Cell Types Analyzed |
| --- | --- | --- | --- | --- |
| Human hematopoiesis | scRNA-seq | 14432 | 12558 | 7 |
| Mouse organogenesis | Temporal scRNA-seq | 158858 | 7798 | 5 |

The count matrix of each dataset normalized with scanpy (*scanpy*.*preprocessing*.*normalize*_*total* and *scanpy*.*preprocessing*.*log*1*p*). For the HHD dataset, we computed pseudotimes using Slingshot [18], with the number of timepoints equal to the depth of the lineage tree computed by Zhang et al. [22]. For the SMO dataset, we use slices from day 11.5, 12.5, and 13.5 embryos. To run the methods on the baseline models in a reasonable amount of compute and training time (24Gb RAM, 24 hours), we performed the following random subsamplings of the HHD data: 2800 cells for SINCERITIES, 2800 cells for GRNBoost2. To run the methods on the baseline models, we performed the following random subsamplings of the SMO data: 12000 cells for ctOTVelo, 12000 cells for OTVelo, 3000 cells for SINCERITIES, 3000 cells for GRNBoost2, 15000 cells for scMTNI.

### 4.2 Benchmarks

We compare the performance of ctOTVelo to state-of-the-art GRN inference models that infer GRNs from transcriptomic data. OTVelo is an optimal transport-based method that infers pseudo-gene velocities to predict GRNs. We train OTVelo using the OT-Velocity package provided by Zhao et al [23]. scMTNI is a GRN inference method that uses cell clusters and a lineage tree to predict GRNs from paired single cell gene expression and ATAC data. scMTNI also has a version that utilizes single cell gene expression only. We train the single cell gene expression-only scMTNI as a baseline for this paper. We use the package scMTNI provided by Zhang et al [22]. SINCERITIES utilizes regularized ridge regression to identify gene regulatory relationships from temporally resolve single cell gene expression data. We trained SINCERITIES using the SINCERITIES package provided by Papili Gao et al[15]. GRNBoost2 is a tree-based GRN inference method that utilizes gradient boosting to identify regulatory relationships from single cell gene expression data. We trained GRNBoost2 using the arboreto package from Moerman et al [12].

### 4.3 Quantitative Evaluation

**Table 2:** Ground truth GRNs.

| Name | Species | Modality | Cell type specific |
| --- | --- | --- | --- |
| UniBind | Human | ChIP-seq | No |
| Cus_ChIP_2 | Human | ChIP-seq | No |
| Cus_KO | Human | Perturbation | No |
| $Cus\_ChIP\_2 \cap Cus\_KO$ | Human | Perturbation + ChIP-seq | No |
| $Cus\_ChIP\_2 \cap UniBind$ | Human | ChIP-seq | No |
| CD34_hematopoietic_stem_cells-derived_proerythroblasts | Human | ChIP-seq | Yes |
| CD34_hematopoietic_stem_cells-derived_proerythroblasts | Human | ChIP-seq | Yes |
| CD14_monocytes | Human | ChIP-seq | Yes |
| CD34_hematopoietic_stem_cells-derived_proerythroblasts | Human | ChIP-seq | Yes |
| Megakaryocytes | Human | ChIP-seq | Yes |
| Erythroid_progenitors | Human | ChIP-seq | Yes |
| R3R4_erythroid_cells | Human | ChIP-seq | Yes |
| GM_B-cells | Human | ChIP-seq | Yes |
| mESC_KD | Mouse | Perturbation | No |
| mESC_chip | Mouse | ChIP-seq | No |
| $mESC\_chip \cap mESC\_KD$ | Mouse | Perturbation + ChIP-seq | No |

To quantitatively evaluate the GRN predictive performance, we two metrics from [22]. For each predicted GRN, we compute the F1 score of the top *k* predicted edges against a ground truth GRN, for *k* ∈ {100, 200, 300, 400, 500, 1000, 2000}. This type of “early” F1 score measures the quality of the top edges in a network, which may be more interpretable for biologists. Finally, we compute the number of predictable transcription factors (TFs), where a predictable TF is a TF in the top *k* predicted edges subgraph of a predicted GRN whose children nodes have a significant overlap with the ground truth target set (via a hypergeometric test with an FDR cutoff of 0.05). This is a stricter measure of recall, where we evaluate the recall of a TF-to-targets subgraph of the GRN.

For ground truth GRNs for the human hematopoiesis dataset, we utilize the ChIP-seq experiments from the UniBind [17] database; ChIP-seq and regulator perturbation experiments from Cusanovich et al. [8]; and the permutations of intersections of ChIP-seq and perturbation networks for a total of five ground truth GRNs. We also divide the UniBind database into cell type specific networks, similar to experiments from Zhang et al [22]. For the ground truth GRNs for the spatiotemporal mouse organogenesis dataset, we utilize the gold standard databases provided in [22]: ChIP-seq experiments from the ENCODE and ESCAPE databases [21, 1], regulator perturbation experiments [14], and the intersection of the edges between the two databases.

We compute F1 and the number of predictable TFs of global (non-cell type specific) predicted GRNs against the five ground truths, and we compute both metrics for the cell type specific ground truths. When evaluating against a global GRN, if a model predicts cell type-specific or lineage/temporal specific GRNs, we evaluate the union of all predicted GRNs against the ground truth. If a model does not predict a cell type specific GRN, we used the predicted global GRN. When evaluating against a cell type-specific GRN, if a model can predict GRNs for a specific cell type, we evaluate using the predicted cell type-specific GRNs. If a model predicts lineage specific GRNs, we evaluate using the predicted lineage-specific GRN derived from cells with a significant composition of that cell type (at least 25%). If a model does not predict a cell type specific GRN, we used the predicted global GRN. We trained ctOTVelo, OTVelo, SINCERITIES, GRNBoost2, and scMTNI on five different seeds. To summarize performance, we calculated the average rank for each metric based on the mean score across the models and datasets for both global and cell type-specific GRN prediction tasks.

### 4.4 Inferring cell type specific GRNs in human hematopoiesis differentiation

We first quantitatively benchmarked the models on the human hematopoiesis differentiation (HHD) dataset. For each metric and ground truth, we computed the ranks of the performances of each model for each value of *k* using the mean and summarized the average rank distribution for the global metrics and the cell type-specific metrics (Figure 2). On the global GRN prediction task, ctOTVelo achieves the strongest performance in F1, with a median average rank of 1.0 and mean average rank of 1.54. In the number of predictable TFs metric, ctOTVelo also has the strongest performance with a median average rank of 1.5 and mean average rank of 1.83. On the cell type specific GRN prediction tasks, ctOTVelo and SINCERITIES have the strongest performance all baselines in the F1 task, with a median rank of 2.0. In the number of predictable TFs metric, ctOTVelo, OTVelo, SINCERITIES, and GRNBoost2 all achieved a median average rank of 3.0, with OTVelo having the highest mean average rank. These results show that ctOTVelo matches or improves overall GRN predictive power for both ground truth and cell type-specific GRNs compared to OTVelo, which excludes cell type specificity, and other baselines. Full quantitative results of each metric are available in Appendix B.

**Figure 2:**
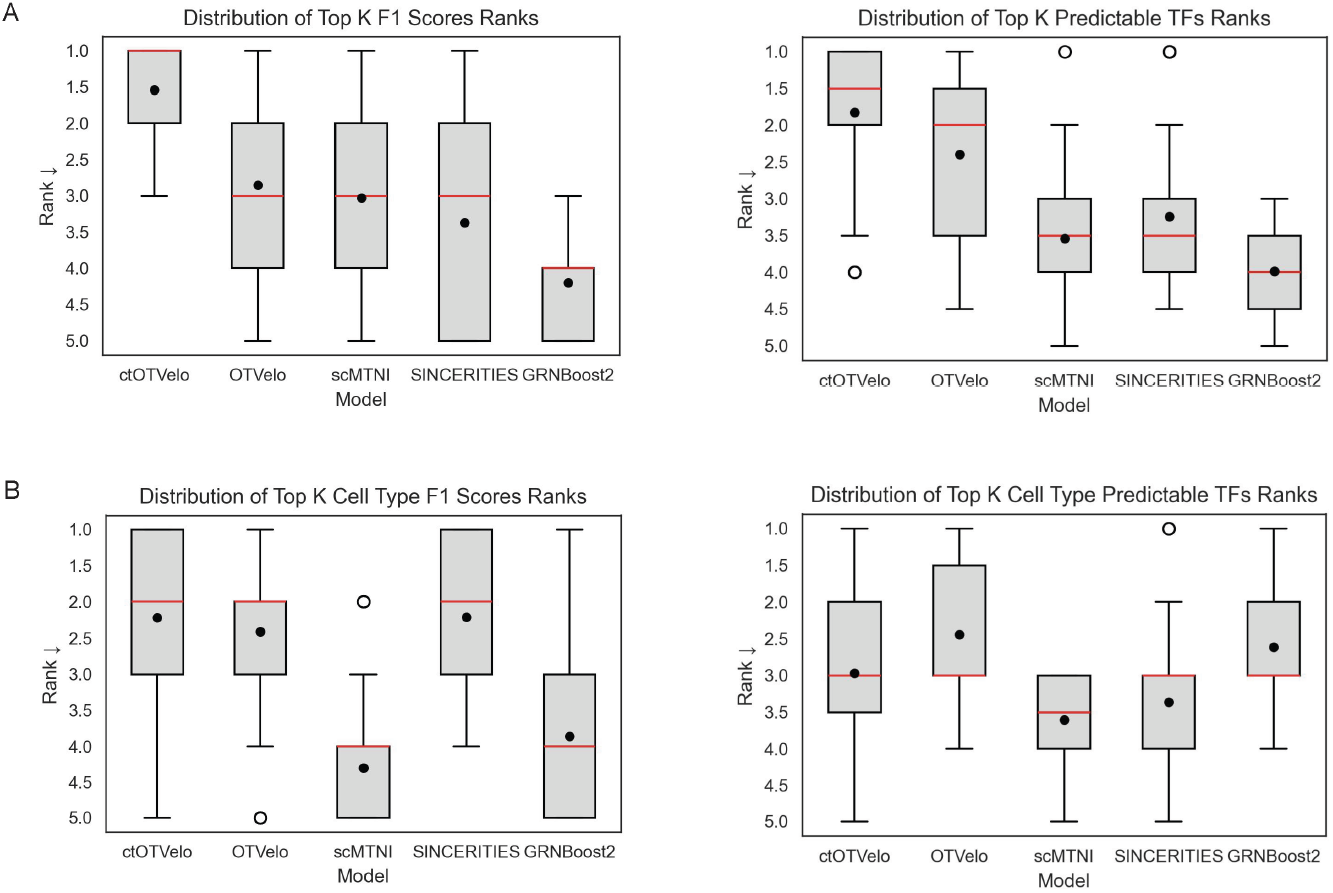
Quantitative performance of GRN inference models on the human hematopoiesis dataset. A. Left: Rank distribution of the models in the top *k* edges F1-score metric performance on the global ground truth GRNs. Right: Rank distribution of the models in the top *k* edges predictable TFs metric performance on the global ground truth GRNs. B. Left: Rank distribution of the models in the top *k* edges F1-score metric performance on the cell type-specific ground truth GRNs. Right: Rank distribution of the models in the top *k* edges predictable TFs metric performance on the cell type-specific ground truth GRNs.

To determine cell type-specific regulatory changes, we examined the cell type graphs from ctOTVelo more closely. We took the pairwise edge set difference between the top 50 predicted edges of each cell type GRN and the top 50 predicted edges of the global GRN (OTVelo), subsetting edges to those found in the respective ground truth cell type GRNs. From these differential GRNs, we observed greater retention of cell type-specific regulatory changes in the predicted cell type specific GRNs compared to the global predicted GRNs (Figure 3). With the hematopoietic stem cell (HSC) cell type, we see that the cell type specific GRNs identifies a greater number of cell type specific regulatory relationships, denoted by a greater number of edges lost in the global GRN compared to number of edges gained. Specifically, *GATA1* is a critical gene in hematopoiesis and modification in GATA1 expression leads to terminal or inhibited erythroid development [10]. Thus, the loss of these *GATA1* relationships in the global GRN in comparison to the HSC-specific GRN shows that ctOTVelo is able to capture HSC-specific regulatory relationships. In the common myeloid progenitor (CMP) cell type, we see that again the global GRN loses a greater number of regulatory edges compared to the cell type specific graph. These edges involved transcription factor *CTCF*, which has been shown to control myeloid cell differentiation through the erythroid lineage [20]. Therefore, the loss of these *CTCF* relationships in the global GRN from the CMP-specific GRN demonstrates ctOTVelo to capture CMP-specific gene regulatory patterns. Overall, ctOTVelo is able to detect cell type specific regulatory patterns that may be missed by global GRNs.

**Figure 3:**
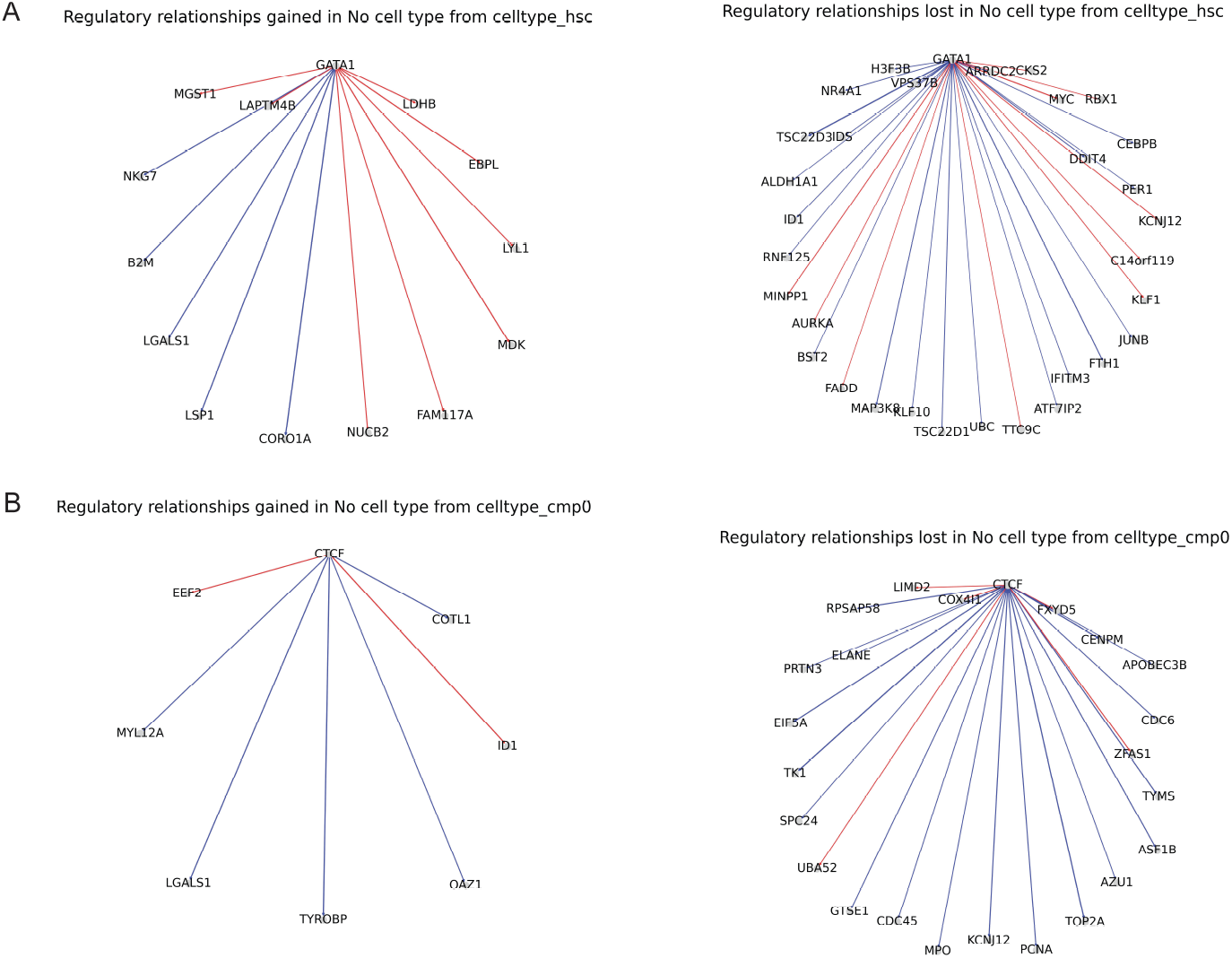
Cell type-specific changes in the gene regulatory networks derived from ctOTVelo. Cell type specific GRN comparison. A. Regulatory relationships in the no cell type GRN gained and lost from the hematopoietic stem cell (HSC) specific GRN. B. Regulatory relationships in the no cell type GRN gained and lost from the common myeloid progenitor (CMP) specific GRN.

### 4.5 Inferring GRNs from spatiotemporal mouse organogenesis

Next we quantitatively benchmarked the models on the spatiotemporal mouse organogenesis (SMO) dataset. For each metric and ground truth, we computed the ranks of the performances of each model (for each value of *k* for the early F1 and predictable TF metrics) using the mean and summarized the average rank distribution for the global metrics (Figure 4. On the global GRN prediction task, ctOTVelo achieves the strongest performance in F1, with a median average rank of 1.0 and mean average rank of 1.61, followed by GRNBoost2, with a median average rank of 2.0 and mean average rank of 2.14. In the number of predictable TFs metric, OTVelo performed the best with a median average rank of 1.0 and mean average rank of 1.43. These results show that cell type specificity matches or improves overall GRN predictive power compared to OTVelo and other baselines. However, OTVelo excels when it comes to detecting the full TF-target sets in the ground truths as seen in the predictable TFs metric. We hypothesize that this can be attributed to cell type specificity focusing on specific TF-target relationships, while the ground truth networks include all experimentally validated relationships. Therefore, a cell type specific model may not retrieve all targets for a TF, which would be desirable behavior. Full quantitative results of each metric are available in Appendix C.

**Figure 4:**
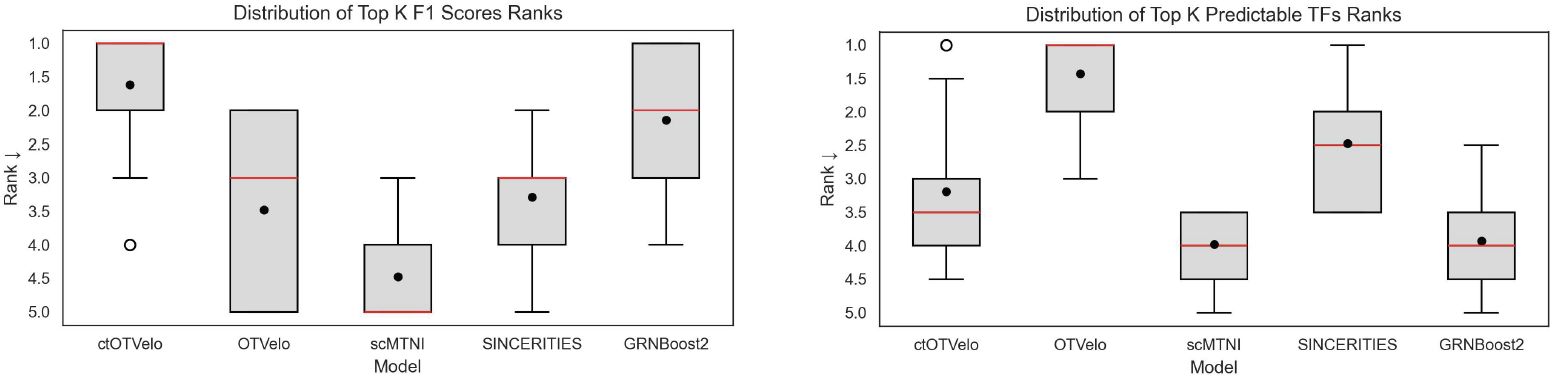
Quantitative performance of GRN inference models on the mouse organogenesis dataset. Left: Rank distribution of the models in the top *k* edges F1-score metric performance on the global ground truth GRNs. Right: Rank distribution of the models in the top *k* edges predictable TFs metric performance on the global ground truth GRNs.

Next we examined the inferred cell type specific GRNs in the same manner as before. We took the pairwise edge set difference between the top 500 predicted edges of each cell type GRN fro ctOTVelo and the top 500 predicted edges of the global GRN (OTVelo), subsetting to edges in the intersection of the ChIP-seq and perturbation gold standard ground truth (Figure 5). In the comparison between the global and liver cell type GRNs, we observe an overall loss in *Tuba1a* and *Jarid2* regulatory relationships in the global GRN compared to the liver cell type GRN. *Tuba1a* has been shown to play a key role in microtubule stabilization (the degradation of which is critical in fibrosis), and *Jarid2* has been linked with embryonic development of the liver [13]. Other studies have shown that *Tuba1a* is also critical for neuronal development [3], and *Jarid2* which temporally regulates neuronal tube development [19] and is crucial in embryonic stem cell differentiation [11], so the mixture of relationships lost and gained is plausible. On the other hand, the global GRN contains regulatory relationships involving *Sox1*, a key marker in neuronal development. The lack of these regulatory relationships in the liver GRN is another supporting sign that the liver-specific GRN is excluding non-liver specific gene regulation. In the comparison between the global and cavity cell type GRNs, we observe an overall gain of regulatory relationships with the *Jarid2* in the cavity GRN. Recent studies by Cho et al. [7] have revealed *Jarid2* as a regulator in proper cardiac development, particularly proper formation of the ventricular wall. The peritoneal and pericardial cavity play a key role in [4] ventricular wall morphogenesis, thus the presence of *Jarid2* regulatory relationships in the cavity-specific GRN is biologically aligned with cavity-specific functions.

**Figure 5:**
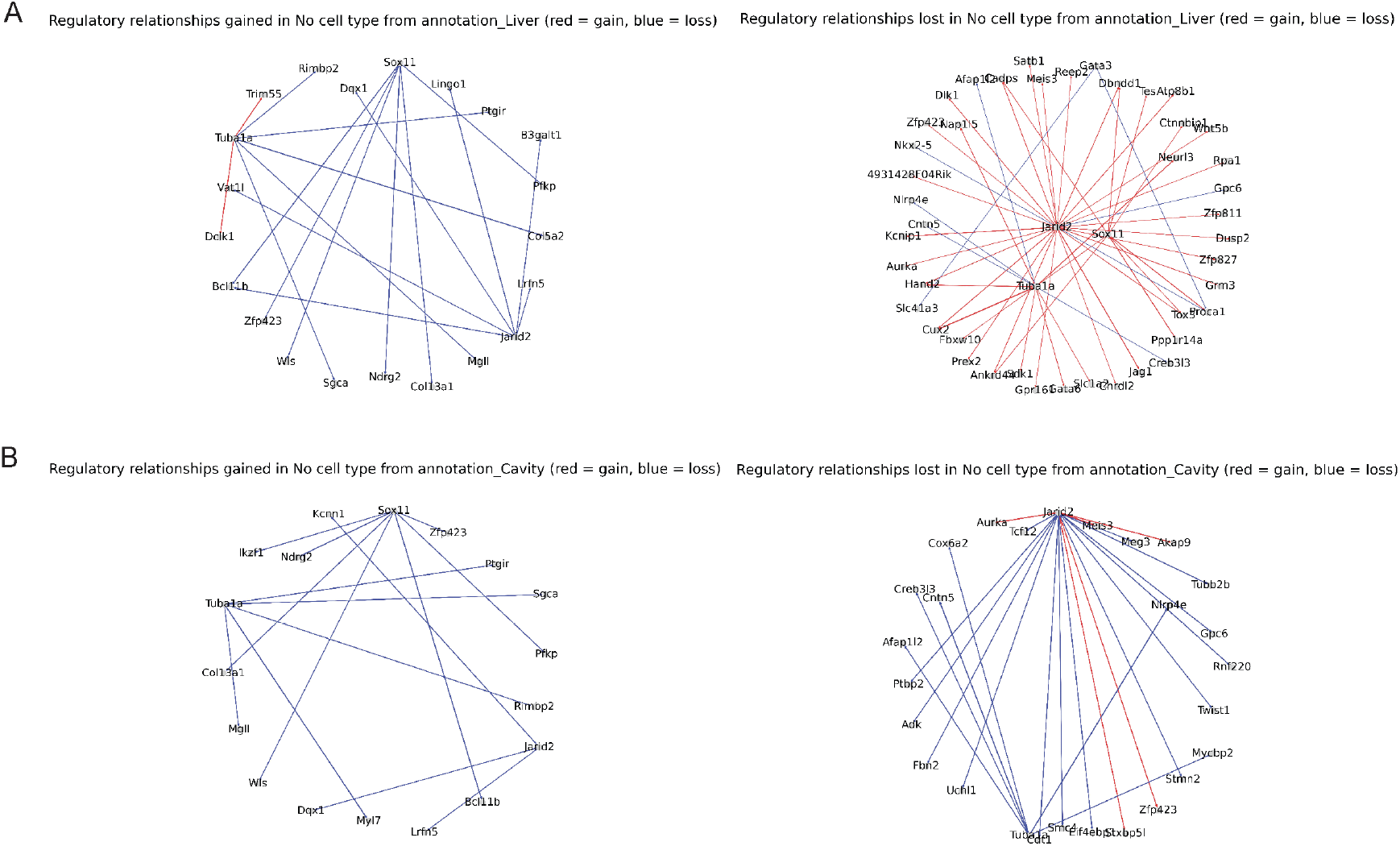
Cell type specific GRN comparison. A. Regulatory relationships gained and lost from the liver specific GRN. B. Regulatory relationships gained and lost from the cavity specific GRN.

## 5 Discussion

In this paper, we present ctOTVelo, an optimal transport based method for inferring cell type specific GRNs. By incorporating cell type information to guide time-lagged correlation, ctOTVelo is able to capture cell type specific gene regulatory relationships. ctOTVelo performs comparably or stronger at GRN prediction compared to state of the art GRN inference models. ctOTVelo’s ability to the generate cell type specific graphs allow for cell type specific downstream analysis.

We also identify some limitations. ctOTVelo’s formulation emphasizes cell type specific interactions, which can lead it to miss global gene regulatory relationships as demonstrated on the mouse organogenesis dataset. Time-lagged correlation is also not a true causal framework, which we leave open to future work. Finally, ctOTvelo requires cell type labels in order to identify cell type specific regulatory interaction - bias that is pressent in the cell type labels will be propagated into the cell type GRN inference, thus ctOTVelo requires robust cell type labeling. ctOTVelo demonstrates the utility of incorporating cell type dynamics into GRN inference for more fine-grained analysis of gene regulatory relationships.

## 6 Code Availability

Code for ctOTVelo and associated analyses presented in this paper is available at https://github.com/seowonchang/ctotvelo.

## Acknowledgments and Disclosure of Funding

SC was a trainee supported under the Brown University Predoctoral Training Program in Biological Data Science (NIH T32 GM149433).

## A Notation

We utilize notation consistent with OTVelo:

**Table S1:** Table of Notations.

| Notation | Description |
| --- | --- |
| $g \in \{1, \dots, m\}$ | Index for genes |
| $t \in \{t_1, \dots, t_N\}$ | Index for time points |
| $n(t) \in \mathbb{R}$ | Number of cells at time $t$ |
| $c_t = \{1, 2, \dots, n(t)\}$ | Index for cell |
| $\mathbf{X} = (\mathbf{X}^{t_1}, \mathbf{X}^{t_2}, \dots, \mathbf{X}^{t_N})$ | Time-stamped gene expression |
| $\mathbf{X}^t \in \mathbb{R}^{c_t \times g}$ | Gene expression at time $t$ |
| $x_g^{c,t} \in \mathbb{R}$ | gene expression of gene $g$ for cell $c$ at time $t$ |
| $D^{t,\tilde{t}} \in \mathbb{R}^{n(t) \times n(\tilde{t})}$ | Difference between cells at time $t$ and $\tilde{t}$ |
| $D_{c,\tilde{c}}^{t,\tilde{t}} \in \mathbb{R}^{n(t) \times n(\tilde{t})}$ | Entry of $D^{t,\tilde{t}}$ representing Euclidean distance between cells $c$ and $\tilde{c}$ |
| $S^t \in \mathbb{R}^{n(t) \times n(t)}$ | Distances between cells at time $t$ of a k-nearest neighbors graph, where $k = \min\{50, 0.2n(t), 0.2n(\tilde{t})\}$ |
| $\ell_{GW} : \mathbb{R}^2 \rightarrow \mathbb{R}^+$ | Function that accounts for misfit pairwise distances |
| $\alpha \in \mathbb{R}$ | Hyperparameter for weighting the Wasserstein and Gromov-Wasserstein terms |
| $T \in \mathbb{R}^{n(t) \times n(t)}$ | Coupling matrix learned from entropic fused Gromov-Wasserstein optimal transport |
| $\Pi(p, q) = \{T : T\mathbf{1} = p, T^T\mathbf{1} = q, T \geq 0\}$ | Family of coupling matrices given source marginal density $p$ and target marginal density $q$ |
| $v_g(x^{t,c}, t) \in \mathbb{R}$ | Velocity of gene $g$ for cell $c$ at time $t$ |

## B Full quantitative metrics for human hematopoiesis

**Table S2:** Mean Top k edges F1 score (± SD) for global GRNs against UniBind.

| Top K edges | ctOTVelo | OTVelo | SINCERITIES | GRNBoost2 | scMTNI |
| --- | --- | --- | --- | --- | --- |
| 100 | **5.12e-04 ± 0.00e+00** | 3.56e-04 ± 0.00e+00 | 3.70e-04 ± 1.70e-04 | 3.14e-04 ± 4.19e-05 | 3.38e-04 ± 4.62e-05 |
| 200 | **1.04e-03 ± 0.00e+00** | 8.41e-04 ± 0.00e+00 | 7.15e-04 ± 3.26e-04 | 6.54e-04 ± 7.83e-05 | 6.89e-04 ± 1.06e-04 |
| 300 | **1.60e-03 ± 0.00e+00** | 1.29e-03 ± 0.00e+00 | 1.06e-03 ± 4.63e-04 | 9.85e-04 ± 1.07e-04 | 1.03e-03 ± 1.28e-04 |
| 400 | **2.11e-03 ± 0.00e+00** | 1.79e-03 ± 0.00e+00 | 1.44e-03 ± 6.20e-04 | 1.32e-03 ± 8.44e-05 | 1.43e-03 ± 1.95e-04 |
| 500 | **2.58e-03 ± 0.00e+00** | 2.19e-03 ± 0.00e+00 | 1.79e-03 ± 7.54e-04 | 1.72e-03 ± 7.50e-05 | 1.80e-03 ± 2.70e-04 |
| 1000 | **4.89e-03 ± 0.00e+00** | 4.60e-03 ± 0.00e+00 | 3.50e-03 ± 1.54e-03 | 3.51e-03 ± 1.41e-04 | 3.55e-03 ± 4.71e-04 |
| 2000 | **9.43e-03 ± 0.00e+00** | 9.28e-03 ± 0.00e+00 | 7.02e-03 ± 1.48e-03 | 6.92e-03 ± 1.91e-04 | 6.64e-03 ± 7.90e-04 |

**Table S3:** Mean Top k edges F1 score (± SD) for global GRNs against Cus_ChIP_2.

| Top K edges | ctOTVelo | OTVelo | SINCERITIES | GRNBoost2 | scMTNI |
| --- | --- | --- | --- | --- | --- |
| 100 | **3.46e-04 ± 0.00e+00** | 1.48e-04 ± 0.00e+00 | 2.92e-04 ± 4.48e-04 | 1.53e-04 ± 2.88e-05 | 2.63e-04 ± 1.19e-04 |
| 200 | **7.65e-04 ± 0.00e+00** | 2.72e-04 ± 0.00e+00 | 5.68e-04 ± 9.27e-04 | 3.50e-04 ± 8.17e-05 | 5.54e-04 ± 1.31e-04 |
| 300 | **1.28e-03 ± 0.00e+00** | 4.44e-04 ± 5.42e-20 | 8.04e-04 ± 1.31e-03 | 5.13e-04 ± 7.86e-05 | 8.35e-04 ± 2.27e-04 |
| 400 | **1.67e-03 ± 2.17e-19** | 6.16e-04 ± 0.00e+00 | 1.06e-03 ± 1.73e-03 | 6.60e-04 ± 8.15e-05 | 1.13e-03 ± 2.87e-04 |
| 500 | **2.09e-03 ± 0.00e+00** | 8.61e-04 ± 0.00e+00 | 1.31e-03 ± 2.10e-03 | 8.16e-04 ± 1.13e-04 | 1.43e-03 ± 3.00e-04 |
| 1000 | **4.13e-03 ± 0.00e+00** | 1.98e-03 ± 0.00e+00 | 2.02e-03 ± 2.99e-03 | 1.94e-03 ± 1.45e-04 | 3.02e-03 ± 4.35e-04 |
| 2000 | **7.99e-03 ± 0.00e+00** | 4.30e-03 ± 0.00e+00 | 3.42e-03 ± 4.09e-03 | 4.10e-03 ± 1.62e-04 | 5.63e-03 ± 6.00e-04 |

**Table S4:** Mean Top k edges F1 score (± SD) for global GRNs against Cus_KO_0.01.

| Top K edges | ctOTVelo | OTVelo | SINCERITIES | GRNBoost2 | scMTNI |
| --- | --- | --- | --- | --- | --- |
| 100 | **8.53e-04 ± 0.00e+00** | 7.31e-04 ± 0.00e+00 | 6.34e-04 ± 2.02e-04 | 6.82e-04 ± 9.75e-05 | 7.33e-04 ± 1.77e-04 |
| 200 | **1.46e-03 ± 0.00e+00** | **1.46e-03 ± 0.00e+00** | 1.13e-03 ± 3.30e-04 | 1.35e-03 ± 1.86e-04 | 1.30e-03 ± 2.63e-04 |
| 300 | 2.12e-03 ± 0.00e+00 | **2.48e-03 ± 0.00e+00** | 1.70e-03 ± 4.13e-04 | 1.95e-03 ± 1.40e-04 | 2.04e-03 ± 3.45e-04 |
| 400 | 2.84e-03 ± 0.00e+00 | **3.62e-03 ± 0.00e+00** | 2.43e-03 ± 6.96e-04 | 2.69e-03 ± 2.49e-04 | 2.70e-03 ± 4.06e-04 |
| 500 | 3.61e-03 ± 0.00e+00 | **4.46e-03 ± 0.00e+00** | 3.01e-03 ± 8.86e-04 | 3.42e-03 ± 2.03e-04 | 3.37e-03 ± 4.20e-04 |
| 1000 | 7.41e-03 ± 0.00e+00 | **8.24e-03 ± 0.00e+00** | 6.22e-03 ± 2.11e-03 | 6.55e-03 ± 4.10e-04 | 6.72e-03 ± 7.06e-04 |
| 2000 | **1.49e-02 ± 1.73e-18** | 1.49e-02 ± 0.00e+00 | 1.17e-02 ± 2.62e-03 | 1.25e-02 ± 3.47e-04 | 1.30e-02 ± 9.45e-04 |

**Table S5:** Mean Top k edges F1 score (± SD) for global GRNs against Cus_KO_0.01_Cus_ChIP_2_intersect.

| Top K edges | ctOTVelo | OTVelo | SINCERITIES | GRNBoost2 | scMTNI |
| --- | --- | --- | --- | --- | --- |
| 100 | **5.38e-03 ± 0.00e+00** | 2.99e-03 ± 0.00e+00 | 2.87e-03 ± 2.66e-03 | 3.59e-03 ± 6.55e-04 | 5.27e-03 ± 1.28e-03 |
| 200 | 8.71e-03 ± 0.00e+00 | 6.97e-03 ± 0.00e+00 | 5.34e-03 ± 5.39e-03 | 6.62e-03 ± 2.53e-03 | **1.01e-02 ± 3.34e-03** |
| 300 | 1.18e-02 ± 0.00e+00 | 1.02e-02 ± 0.00e+00 | 9.03e-03 ± 9.33e-03 | 9.37e-03 ± 3.36e-03 | **1.28e-02 ± 3.29e-03** |
| 400 | 1.48e-02 ± 0.00e+00 | 1.15e-02 ± 0.00e+00 | 1.21e-02 ± 1.20e-02 | 1.22e-02 ± 2.58e-03 | **1.64e-02 ± 4.43e-03** |
| 500 | 1.76e-02 ± 0.00e+00 | 1.23e-02 ± 0.00e+00 | 1.51e-02 ± 1.32e-02 | 1.34e-02 ± 2.25e-03 | **1.96e-02 ± 3.57e-03** |
| 1000 | 2.59e-02 ± 3.47e-18 | 1.65e-02 ± 0.00e+00 | **3.45e-02 ± 1.70e-02** | 1.34e-02 ± 2.25e-03 | 3.18e-02 ± 4.54e-03 |
| 2000 | 3.55e-02 ± 0.00e+00 | 2.55e-02 ± 0.00e+00 | **5.53e-02 ± 1.83e-02** | 1.34e-02 ± 2.25e-03 | 5.08e-02 ± 3.03e-03 |

**Table S6:** Mean Top k edges F1 score (± SD) for global GRNs against Cus_KO_0.01_union_intersect.

| Top K edges | ctOTVelo | OTVelo | SINCERITIES | GRNBoost2 | scMTNI |
| --- | --- | --- | --- | --- | --- |
| 100 | **1.21e-02 ± 0.00e+00** | **1.21e-02 ± 0.00e+00** | 9.21e-03 ± 5.54e-03 | 1.03e-02 ± 1.63e-03 | 6.06e-03 ± 9.58e-04 |
| 200 | 2.26e-02 ± 0.00e+00 | **2.40e-02 ± 0.00e+00** | 1.67e-02 ± 1.11e-02 | 1.95e-02 ± 2.66e-03 | 1.27e-02 ± 2.79e-03 |
| 300 | **2.95e-02 ± 3.47e-18** | 2.67e-02 ± 3.47e-18 | 2.70e-02 ± 1.92e-02 | 2.09e-02 ± 3.74e-03 | 1.96e-02 ± 2.33e-03 |
| 400 | **3.46e-02 ± 0.00e+00** | 3.12e-02 ± 3.47e-18 | 3.35e-02 ± 2.33e-02 | 2.09e-02 ± 3.74e-03 | 2.49e-02 ± 2.22e-03 |
| 500 | 3.94e-02 ± 0.00e+00 | 3.87e-02 ± 0.00e+00 | **3.99e-02 ± 2.90e-02** | 2.09e-02 ± 3.74e-03 | 2.92e-02 ± 2.26e-03 |
| 1000 | 6.09e-02 ± 6.94e-18 | 6.26e-02 ± 0.00e+00 | **6.38e-02 ± 3.02e-02** | 2.09e-02 ± 3.74e-03 | 4.88e-02 ± 3.73e-03 |
| 2000 | 9.19e-02 ± 0.00e+00 | 9.28e-02 ± 0.00e+00 | **9.63e-02 ± 2.79e-02** | 2.09e-02 ± 3.74e-03 | 4.90e-02 ± 3.60e-03 |

**Table S7:**
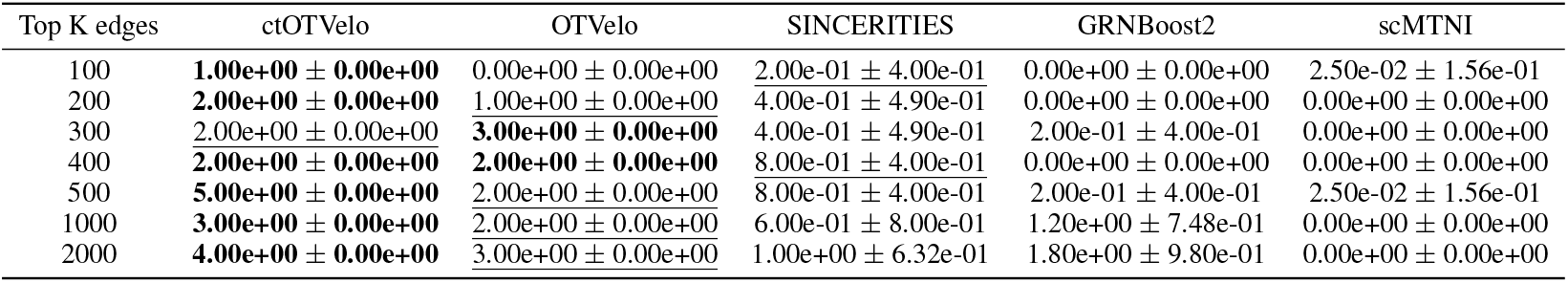
Mean Top k edges Number of predictable TFs (± SD) for global GRNs against UniBind.

**Table S8:** Mean Top k edges Number of predictable TFs (± SD) for global GRNs against Cus_ChIP_2.

| Top K edges | ctOTVelo | OTVelo | SINCERITIES | GRNBoost2 | scMTNI |
| --- | --- | --- | --- | --- | --- |
| 100 | **0.00e+00 ± 0.00e+00** | **0.00e+00 ± 0.00e+00** | **0.00e+00 ± 0.00e+00** | **0.00e+00 ± 0.00e+00** | **0.00e+00 ± 0.00e+00** |
| 200 | **1.00e+00 ± 0.00e+00** | 0.00e+00 ± 0.00e+00 | 0.00e+00 ± 0.00e+00 | 0.00e+00 ± 0.00e+00 | 0.00e+00 ± 0.00e+00 |
| 300 | **2.00e+00 ± 0.00e+00** | 0.00e+00 ± 0.00e+00 | 0.00e+00 ± 0.00e+00 | 0.00e+00 ± 0.00e+00 | 0.00e+00 ± 0.00e+00 |
| 400 | **1.00e+00 ± 0.00e+00** | 0.00e+00 ± 0.00e+00 | 0.00e+00 ± 0.00e+00 | 0.00e+00 ± 0.00e+00 | 0.00e+00 ± 0.00e+00 |
| 500 | **1.00e+00 ± 0.00e+00** | 0.00e+00 ± 0.00e+00 | 0.00e+00 ± 0.00e+00 | 0.00e+00 ± 0.00e+00 | 0.00e+00 ± 0.00e+00 |
| 1000 | **4.00e+00 ± 0.00e+00** | 3.00e+00 ± 0.00e+00 | 0.00e+00 ± 0.00e+00 | 0.00e+00 ± 0.00e+00 | 0.00e+00 ± 0.00e+00 |
| 2000 | **7.00e+00 ± 0.00e+00** | 2.00e+00 ± 0.00e+00 | 4.00e-01 ± 4.90e-01 | 0.00e+00 ± 0.00e+00 | 2.50e-02 ± 1.56e-01 |

**Table S9:** Mean Top k edges Number of predictable TFs (± SD) for global GRNs against Cus_KO_0.01.

| Top K edges | ctOTVelo | OTVelo | SINCERITIES | GRNBoost2 | scMTNI |
| --- | --- | --- | --- | --- | --- |
| 100 | 0.00e+00 ± 0.00e+00 | 0.00e+00 ± 0.00e+00 | **2.00e-01 ± 4.00e-01** | 0.00e+00 ± 0.00e+00 | 5.00e-02 ± 3.12e-01 |
| 200 | 0.00e+00 ± 0.00e+00 | 0.00e+00 ± 0.00e+00 | **2.00e-01 ± 4.00e-01** | 0.00e+00 ± 0.00e+00 | 5.00e-02 ± 2.18e-01 |
| 300 | **1.00e+00 ± 0.00e+00** | 0.00e+00 ± 0.00e+00 | 0.00e+00 ± 0.00e+00 | 0.00e+00 ± 0.00e+00 | <u>2.50e-02 ± 1.56e-01</u> |
| 400 | **1.00e+00 ± 0.00e+00** | **1.00e+00 ± 0.00e+00** | 0.00e+00 ± 0.00e+00 | 0.00e+00 ± 0.00e+00 | <u>7.50e-02 ± 2.63e-01</u> |
| 500 | **1.00e+00 ± 0.00e+00** | **1.00e+00 ± 0.00e+00** | 2.00e-01 ± 4.00e-01 | 0.00e+00 ± 0.00e+00 | 7.50e-02 ± 2.63e-01 |
| 1000 | **2.00e+00 ± 0.00e+00** | **2.00e+00 ± 0.00e+00** | <u>2.00e-01 ± 4.00e-01</u> | 0.00e+00 ± 0.00e+00 | 5.00e-02 ± 2.18e-01 |
| 2000 | **7.00e+00 ± 0.00e+00** | <u>3.00e+00 ± 0.00e+00</u> | 0.00e+00 ± 0.00e+00 | 0.00e+00 ± 0.00e+00 | 5.00e-02 ± 2.18e-01 |

**Table S10:** Mean Top k edges Number of predictable TFs (± SD) for global GRNs against Cus_KO_0.01_Cus_ChIP_2_intersect.

| Top K edges | ctOTVelo | OTVelo | SINCERITIES | GRNBoost2 | scMTNI |
| --- | --- | --- | --- | --- | --- |
| 100 | **1.00e+00 ± 0.00e+00** | 0.00e+00 ± 0.00e+00 | 0.00e+00 ± 0.00e+00 | 0.00e+00 ± 0.00e+00 | 0.00e+00 ± 0.00e+00 |
| 200 | **1.00e+00 ± 0.00e+00** | **1.00e+00 ± 0.00e+00** | 0.00e+00 ± 0.00e+00 | 0.00e+00 ± 0.00e+00 | 0.00e+00 ± 0.00e+00 |
| 300 | **1.00e+00 ± 0.00e+00** | **1.00e+00 ± 0.00e+00** | 0.00e+00 ± 0.00e+00 | 0.00e+00 ± 0.00e+00 | 0.00e+00 ± 0.00e+00 |
| 400 | **1.00e+00 ± 0.00e+00** | **1.00e+00 ± 0.00e+00** | <u>2.00e-01 ± 4.00e-01</u> | 0.00e+00 ± 0.00e+00 | 0.00e+00 ± 0.00e+00 |
| 500 | **1.00e+00 ± 0.00e+00** | **1.00e+00 ± 0.00e+00** | 4.00e-01 ± 4.90e-01 | 0.00e+00 ± 0.00e+00 | 0.00e+00 ± 0.00e+00 |
| 1000 | 2.00e+00 ± 0.00e+00 | **3.00e+00 ± 0.00e+00** | 0.00e+00 ± 0.00e+00 | 0.00e+00 ± 0.00e+00 | 0.00e+00 ± 0.00e+00 |
| 2000 | <u>1.00e+00 ± 0.00e+00</u> | **2.00e+00 ± 0.00e+00** | 0.00e+00 ± 0.00e+00 | 0.00e+00 ± 0.00e+00 | 0.00e+00 ± 0.00e+00 |

**Table S11:** Mean Top k edges Number of predictable TFs (± SD) for global GRNs against Cus_KO_0.01_union_intersect.

| Top K edges | ctOTVelo | OTVelo | SINCERITIES | GRNBoost2 | scMTNI |
| --- | --- | --- | --- | --- | --- |
| 100 | 0.00e+00 ± 0.00e+00 | 0.00e+00 ± 0.00e+00 | **2.00e-01 ± 4.00e-01** | 0.00e+00 ± 0.00e+00 | 0.00e+00 ± 0.00e+00 |
| 200 | 0.00e+00 ± 0.00e+00 | **1.00e+00 ± 0.00e+00** | 0.00e+00 ± 0.00e+00 | 0.00e+00 ± 0.00e+00 | 0.00e+00 ± 0.00e+00 |
| 300 | **1.00e+00 ± 0.00e+00** | 0.00e+00 ± 0.00e+00 | 0.00e+00 ± 0.00e+00 | 0.00e+00 ± 0.00e+00 | 0.00e+00 ± 0.00e+00 |
| 400 | 0.00e+00 ± 0.00e+00 | 0.00e+00 ± 0.00e+00 | 0.00e+00 ± 0.00e+00 | 0.00e+00 ± 0.00e+00 | **2.50e-02 ± 1.56e-01** |
| 500 | 0.00e+00 ± 0.00e+00 | 0.00e+00 ± 0.00e+00 | **2.00e-01 ± 4.00e-01** | 0.00e+00 ± 0.00e+00 | <u>2.50e-02 ± 1.56e-01</u> |
| 1000 | **1.00e+00 ± 0.00e+00** | **1.00e+00 ± 0.00e+00** | 0.00e+00 ± 0.00e+00 | 0.00e+00 ± 0.00e+00 | <u>2.50e-02 ± 1.56e-01</u> |
| 2000 | 0.00e+00 ± 0.00e+00 | **2.00e+00 ± 0.00e+00** | 0.00e+00 ± 0.00e+00 | 0.00e+00 ± 0.00e+00 | <u>2.50e-02 ± 1.56e-01</u> |

**Table S12:** Mean Top k edges F1 score (± SD) for Hematopoietic stem cells GRNs against CD34_hematopoietic_stem_cells_CD34CD133_hematopoietic_progenitors.

| Top K edges | ctOTVelo | OTVelo | SINCERITIES | GRNBoost2 | scMTNI |
| --- | --- | --- | --- | --- | --- |
| 100 | 3.80e-03 ± 0.00e+00 | 3.96e-03 ± 0.00e+00 | **4.29e-03 ± 1.53e-03** | 3.94e-03 ± 4.65e-04 | 4.24e-03 ± 9.72e-04 |
| 200 | 7.12e-03 ± 8.67e-19 | 6.97e-03 ± 8.67e-19 | **7.73e-03 ± 3.54e-03** | 7.67e-03 ± 4.48e-04 | 7.41e-03 ± 3.22e-03 |
| 300 | 1.06e-02 ± 0.00e+00 | 1.02e-02 ± 0.00e+00 | **1.12e-02 ± 5.11e-03** | 1.12e-02 ± 5.35e-04 | 8.09e-03 ± 3.77e-03 |
| 400 | 1.46e-02 ± 1.73e-18 | 1.38e-02 ± 0.00e+00 | **1.50e-02 ± 6.55e-03** | 1.48e-02 ± 8.04e-04 | 8.09e-03 ± 3.77e-03 |
| 500 | 1.82e-02 ± 0.00e+00 | 1.73e-02 ± 0.00e+00 | <u>1.88e-02 ± 8.26e-03</u> | **1.92e-02 ± 9.43e-04** | 8.09e-03 ± 3.77e-03 |
| 1000 | 3.47e-02 ± 0.00e+00 | 3.24e-02 ± 0.00e+00 | **3.81e-02 ± 1.57e-02** | 2.09e-02 ± 1.40e-03 | 8.09e-03 ± 3.77e-03 |
| 2000 | <u>7.01e-02 ± 0.00e+00</u> | 6.19e-02 ± 0.00e+00 | **7.83e-02 ± 2.23e-02** | 2.09e-02 ± 1.40e-03 | 8.09e-03 ± 3.77e-03 |

**Table S13:** Mean Top k edges F1 score (± SD) for Hematopoietic stem cells GRNs against CD34_hematopoietic_stem_cells-derived_proerythroblasts.

| Top K edges | ctOTVelo | OTVelo | SINCERITIES | GRNBoost2 | scMTNI |
| --- | --- | --- | --- | --- | --- |
| 100 | **2.67e-02 ± 0.00e+00** | **2.67e-02 ± 0.00e+00** | 1.70e-02 ± 1.08e-02 | <u>2.37e-02 ± 5.98e-04</u> | 7.21e-03 ± 4.49e-03 |
| 200 | **5.35e-02 ± 6.94e-18** | 5.32e-02 ± 0.00e+00 | 3.43e-02 ± 2.14e-02 | 4.47e-02 ± 1.40e-03 | 9.55e-03 ± 8.86e-03 |
| 300 | **7.84e-02 ± 0.00e+00** | 7.81e-02 ± 0.00e+00 | 5.13e-02 ± 3.01e-02 | 6.55e-02 ± 1.97e-03 | 1.08e-02 ± 1.16e-02 |
| 400 | **1.03e-01 ± 0.00e+00** | <u>1.02e-01 ± 1.39e-17</u> | 6.74e-02 ± 3.94e-02 | 6.91e-02 ± 4.18e-03 | 1.13e-02 ± 1.25e-02 |
| 500 | **1.26e-01 ± 0.00e+00** | 1.25e-01 ± 0.00e+00 | 8.24e-02 ± 4.60e-02 | 6.91e-02 ± 4.18e-03 | 1.17e-02 ± 1.33e-02 |
| 1000 | **2.32e-01 ± 0.00e+00** | <u>2.31e-01 ± 0.00e+00</u> | 1.47e-01 ± 6.50e-02 | 6.91e-02 ± 4.18e-03 | 1.18e-02 ± 1.34e-02 |
| 2000 | **4.04e-01 ± 5.55e-17** | <u>3.96e-01 ± 0.00e+00</u> | 2.11e-01 ± 7.61e-02 | 6.91e-02 ± 4.18e-03 | 1.18e-02 ± 1.34e-02 |

**Table S14:**
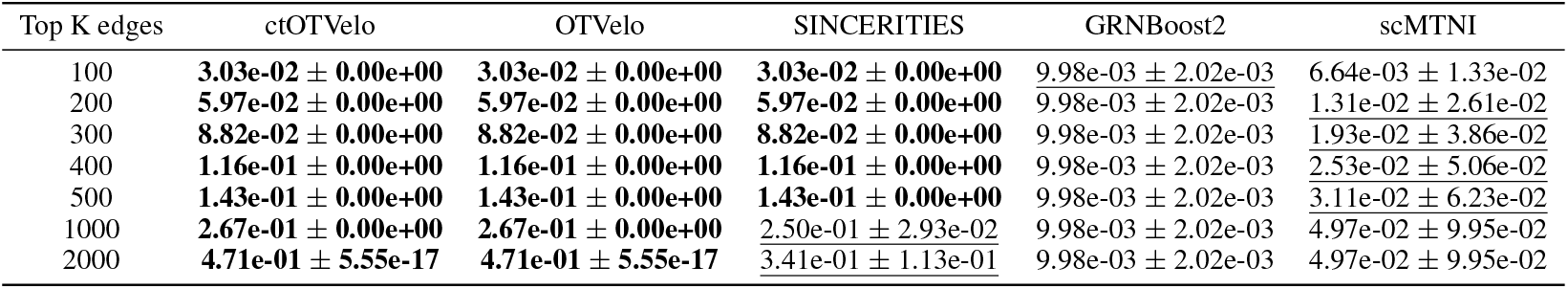
Mean Top k edges F1 score (± SD) for Monocyte GRNs against CD14_monocytes.

| Top K edges | ctOTVelo | OTVelo | SINCERITIES | GRNBoost2 | scMTNI |
| --- | --- | --- | --- | --- | --- |
| 100 | **3.03e-02** ± **0.00e+00** | **3.03e-02** ± **0.00e+00** | **3.03e-02** ± **0.00e+00** | 9.98e-03 ± 2.02e-03 | 6.64e-03 ± 1.33e-02 |
| 200 | **5.97e-02** ± **0.00e+00** | **5.97e-02** ± **0.00e+00** | **5.97e-02** ± **0.00e+00** | 9.98e-03 ± 2.02e-03 | <u>1.31e-02</u> ± <u>2.61e-02</u> |
| 300 | **8.82e-02** ± **0.00e+00** | **8.82e-02** ± **0.00e+00** | **8.82e-02** ± **0.00e+00** | 9.98e-03 ± 2.02e-03 | <u>1.93e-02</u> ± <u>3.86e-02</u> |
| 400 | **1.16e-01** ± **0.00e+00** | **1.16e-01** ± **0.00e+00** | **1.16e-01** ± **0.00e+00** | 9.98e-03 ± 2.02e-03 | <u>2.53e-02</u> ± <u>5.06e-02</u> |
| 500 | **1.43e-01** ± **0.00e+00** | **1.43e-01** ± **0.00e+00** | **1.43e-01** ± **0.00e+00** | 9.98e-03 ± 2.02e-03 | <u>3.11e-02</u> ± <u>6.23e-02</u> |
| 1000 | **2.67e-01** ± **0.00e+00** | **2.67e-01** ± **0.00e+00** | <u>2.50e-01</u> ± <u>2.93e-02</u> | 9.98e-03 ± 2.02e-03 | <u>4.97e-02</u> ± <u>9.95e-02</u> |
| 2000 | **4.71e-01** ± **5.55e-17** | **4.71e-01** ± **5.55e-17** | <u>3.41e-01</u> ± <u>1.13e-01</u> | 9.98e-03 ± 2.02e-03 | <u>4.97e-02</u> ± <u>9.95e-02</u> |

**Table S15:** Mean Top k edges F1 score (± SD) for Common myeloid progenitor GRNs against megakaryocytes.

| Top K edges | ctOTVelo | OTVelo | SINCERITIES | GRNBoost2 | scMTNI |
| --- | --- | --- | --- | --- | --- |
| 100 | 4.38e-03 ± 1.11e-03 | 5.42e-03 ± 0.00e+00 | 5.90e-03 ± 3.11e-03 | **6.50e-03** ± **2.57e-04** | 6.03e-03 ± 1.74e-03 |
| 200 | 8.25e-03 ± 9.91e-04 | 8.47e-03 ± 0.00e+00 | **1.21e-02** ± **6.75e-03** | <u>1.19e-02</u> ± <u>5.98e-04</u> | 1.10e-02 ± 4.57e-03 |
| 300 | 1.22e-02 ± 3.93e-04 | 1.14e-02 ± 0.00e+00 | **1.87e-02** ± **9.75e-03** | <u>1.62e-02</u> ± <u>4.83e-04</u> | 1.57e-02 ± 7.38e-03 |
| 400 | 1.61e-02 ± 8.28e-04 | 1.46e-02 ± 1.73e-18 | **2.53e-02** ± **1.25e-02** | <u>2.01e-02</u> ± <u>2.31e-04</u> | 1.78e-02 ± 8.39e-03 |
| 500 | 2.03e-02 ± 1.55e-03 | 1.82e-02 ± 0.00e+00 | **3.25e-02** ± **1.48e-02** | <u>2.46e-02</u> ± <u>7.20e-04</u> | 1.84e-02 ± 8.77e-03 |
| 1000 | <u>4.19e-02</u> ± <u>6.17e-03</u> | 3.41e-02 ± 0.00e+00 | **6.81e-02** ± **2.65e-02** | 3.48e-02 ± 1.59e-03 | 1.84e-02 ± 8.77e-03 |
| 2000 | <u>8.59e-02</u> ± <u>1.85e-02</u> | 6.08e-02 ± 6.94e-18 | **1.25e-01** ± **4.73e-02** | 3.48e-02 ± 1.59e-03 | 1.84e-02 ± 8.77e-03 |

**Table S16:** Mean Top k edges F1 score (± SD) for Common myeloid progenitor GRNs against erythroid_progenitors.

| Top K edges | ctOTVelo | OTVelo | SINCERITIES | GRNBoost2 | scMTNI |
| --- | --- | --- | --- | --- | --- |
| 100 | **2.49e-02** ± **6.94e-18** | 2.49e-02 ± 0.00e+00 | <u>2.49e-02</u> ± <u>0.00e+00</u> | 9.54e-03 ± 1.67e-03 | 2.21e-02 ± 1.03e-02 |
| 200 | **4.92e-02** ± **1.39e-17** | <u>4.92e-02</u> ± <u>0.00e+00</u> | <u>4.92e-02</u> ± <u>0.00e+00</u> | 9.54e-03 ± 1.67e-03 | 2.42e-02 ± 1.22e-02 |
| 300 | <u>7.29e-02</u> ± <u>1.39e-17</u> | **7.29e-02** ± **0.00e+00** | **7.29e-02** ± **0.00e+00** | 9.54e-03 ± 1.67e-03 | 2.42e-02 ± 1.22e-02 |
| 400 | **9.61e-02** ± **1.39e-17** | <u>9.61e-02</u> ± <u>0.00e+00</u> | <u>9.61e-02</u> ± <u>0.00e+00</u> | 9.54e-03 ± 1.67e-03 | 2.42e-02 ± 1.22e-02 |
| 500 | <u>1.19e-01</u> ± <u>0.00e+00</u> | **1.19e-01** ± **1.39e-17** | **1.19e-01** ± **1.39e-17** | 9.54e-03 ± 1.67e-03 | 2.42e-02 ± 1.22e-02 |
| 1000 | <u>2.24e-01</u> ± <u>2.78e-17</u> | **2.24e-01** ± **0.00e+00** | 2.18e-01 ± 1.17e-02 | 9.54e-03 ± 1.67e-03 | 2.42e-02 ± 1.22e-02 |
| 2000 | <u>4.03e-01</u> ± <u>0.00e+00</u> | **4.03e-01** ± **5.55e-17** | 3.13e-01 ± 8.18e-02 | 9.54e-03 ± 1.67e-03 | 2.42e-02 ± 1.22e-02 |

**Table S17:** Mean Top k edges F1 score (± SD) for Common myeloid progenitor GRNs against B-cells.

| Top K edges | ctOTVelo | OTVelo | SINCERITIES | GRNBoost2 | scMTNI |
| --- | --- | --- | --- | --- | --- |
| 100 | **2.81e-02** ± **6.94e-18** | 2.81e-02 ± 0.00e+00 | <u>2.81e-02</u> ± <u>0.00e+00</u> | 9.59e-03 ± 1.74e-03 | 2.31e-02 ± 1.09e-02 |
| 200 | **5.55e-02** ± **1.39e-17** | <u>5.55e-02</u> ± <u>0.00e+00</u> | <u>5.55e-02</u> ± <u>0.00e+00</u> | 9.59e-03 ± 1.74e-03 | 2.42e-02 ± 1.22e-02 |
| 300 | **8.20e-02** ± **2.78e-17** | <u>8.20e-02</u> ± <u>0.00e+00</u> | <u>8.20e-02</u> ± <u>0.00e+00</u> | 9.59e-03 ± 1.74e-03 | 2.42e-02 ± 1.22e-02 |
| 400 | **1.08e-01** ± **4.16e-17** | <u>1.08e-01</u> ± <u>1.39e-17</u> | <u>1.08e-01</u> ± <u>1.39e-17</u> | 9.59e-03 ± 1.74e-03 | 2.42e-02 ± 1.22e-02 |
| 500 | **1.33e-01** ± **0.00e+00** | **1.33e-01** ± **0.00e+00** | **1.33e-01** ± **0.00e+00** | 9.59e-03 ± 1.74e-03 | 2.42e-02 ± 1.22e-02 |
| 1000 | **2.50e-01** ± **2.78e-17** | <u>2.50e-01</u> ± <u>0.00e+00</u> | 2.38e-01 ± 2.25e-02 | 9.59e-03 ± 1.74e-03 | 2.42e-02 ± 1.22e-02 |
| 2000 | <u>4.44e-01</u> ± <u>0.00e+00</u> | **4.44e-01** ± **5.55e-17** | 3.30e-01 ± 1.01e-01 | 9.59e-03 ± 1.74e-03 | 2.42e-02 ± 1.22e-02 |

**Table S18:** Mean Top k edges F1 score (± SD) for Common myeloid progenitor GRNs against R3R4_erythroid_cells.

| Top K edges | ctOTVelo | OTVelo | SINCERITIES | GRNBoost2 | scMTNI |
| --- | --- | --- | --- | --- | --- |
| 100 | **2.33e-02** ± **0.00e+00** | **2.33e-02** ± **0.00e+00** | **2.33e-02** ± **0.00e+00** | 1.81e-02 ± 9.90e-04 | <u>2.04e-02</u> ± <u>1.03e-02</u> |
| 200 | **4.61e-02** ± **6.94e-18** | <u>4.61e-02</u> ± <u>0.00e+00</u> | <u>4.61e-02</u> ± <u>0.00e+00</u> | 1.81e-02 ± 9.90e-04 | 2.48e-02 ± 1.44e-02 |
| 300 | <u>6.84e-02</u> ± <u>2.78e-17</u> | **6.84e-02** ± **0.00e+00** | **6.84e-02** ± **0.00e+00** | 1.81e-02 ± 9.90e-04 | 2.48e-02 ± 1.44e-02 |
| 400 | **9.02e-02** ± **2.78e-17** | <u>9.02e-02</u> ± <u>0.00e+00</u> | <u>9.02e-02</u> ± <u>0.00e+00</u> | 1.81e-02 ± 9.90e-04 | 2.48e-02 ± 1.44e-02 |
| 500 | **1.11e-01** ± **0.00e+00** | **1.11e-01** ± **0.00e+00** | **1.11e-01** ± **0.00e+00** | 1.81e-02 ± 9.90e-04 | <u>2.48e-02</u> ± <u>1.44e-02</u> |
| 1000 | **2.11e-01** ± **8.33e-17** | <u>2.11e-01</u> ± <u>2.78e-17</u> | 1.89e-01 ± 2.99e-02 | 1.81e-02 ± 9.90e-04 | 2.48e-02 ± 1.44e-02 |
| 2000 | <u>3.82e-01</u> ± <u>1.11e-16</u> | **3.82e-01** ± **0.00e+00** | 2.92e-01 ± 1.11e-01 | 1.81e-02 ± 9.90e-04 | 2.48e-02 ± 1.44e-02 |

**Table S19:** Mean Top k edges F1 score (± SD) for Granulocyte-marophage progenitors GRNs against GM_B-cells.

| Top K edges | ctOTVelo | OTVelo | SINCERITIES | GRNBoost2 | scMTNI |
| --- | --- | --- | --- | --- | --- |
| 100 | 2.59e-04 ± 0.00e+00 | 3.73e-04 ± 0.00e+00 | **4.97e-04 ± 2.67e-04** | 3.48e-04 ± 4.37e-05 | 2.75e-04 ± 1.46e-04 |
| 200 | 6.31e-04 ± 0.00e+00 | 8.80e-04 ± 1.08e-19 | **1.00e-03 ± 4.94e-04** | 7.02e-04 ± 8.94e-05 | 6.11e-04 ± 2.64e-04 |
| 300 | 1.12e-03 ± 0.00e+00 | 1.43e-03 ± 0.00e+00 | **1.46e-03 ± 6.91e-04** | 1.04e-03 ± 8.10e-05 | 1.02e-03 ± 3.10e-04 |
| 400 | 1.58e-03 ± 2.17e-19 | **2.09e-03 ± 0.00e+00** | 1.84e-03 ± 8.44e-04 | 1.45e-03 ± 6.69e-05 | 1.43e-03 ± 3.40e-04 |
| 500 | 2.07e-03 ± 0.00e+00 | **2.62e-03 ± 0.00e+00** | 2.20e-03 ± 9.77e-04 | 1.80e-03 ± 5.97e-05 | 1.82e-03 ± 3.93e-04 |
| 1000 | 4.38e-03 ± 0.00e+00 | **5.45e-03 ± 0.00e+00** | 4.17e-03 ± 1.65e-03 | 3.72e-03 ± 1.08e-04 | 3.97e-03 ± 5.54e-04 |
| 2000 | 9.50e-03 ± 0.00e+00 | **1.10e-02 ± 0.00e+00** | 9.06e-03 ± 2.28e-03 | 7.63e-03 ± 1.56e-04 | 7.12e-03 ± 2.05e-03 |

**Table S20:** Mean Top k edges F1 score (± SD) for CD34 GRNs against CD34_hematopoietic_stem_cells_CD34CD133_hematopoietic_progenitors.

| Top K edges | ctOTVelo | OTVelo | SINCERITIES | GRNBoost2 | scMTNI |
| --- | --- | --- | --- | --- | --- |
| 100 | 3.96e-03 ± 0.00e+00 | 3.96e-03 ± 0.00e+00 | **4.29e-03 ± 1.53e-03** | 3.94e-03 ± 4.65e-04 | 4.24e-03 ± 9.72e-04 |
| 200 | 7.66e-03 ± 0.00e+00 | 6.97e-03 ± 8.67e-19 | **7.73e-03 ± 3.54e-03** | 7.67e-03 ± 4.48e-04 | 7.41e-03 ± 3.22e-03 |
| 300 | 1.06e-02 ± 0.00e+00 | 1.02e-02 ± 0.00e+00 | **1.12e-02 ± 5.11e-03** | 1.12e-02 ± 5.35e-04 | 8.09e-03 ± 3.77e-03 |
| 400 | 1.36e-02 ± 0.00e+00 | 1.38e-02 ± 0.00e+00 | **1.50e-02 ± 6.55e-03** | 1.48e-02 ± 8.04e-04 | 8.09e-03 ± 3.77e-03 |
| 500 | 1.69e-02 ± 0.00e+00 | 1.73e-02 ± 0.00e+00 | 1.88e-02 ± 8.26e-03 | **1.92e-02 ± 9.43e-04** | 8.09e-03 ± 3.77e-03 |
| 1000 | 3.13e-02 ± 0.00e+00 | 3.24e-02 ± 0.00e+00 | **3.81e-02 ± 1.57e-02** | 2.09e-02 ± 1.40e-03 | 8.09e-03 ± 3.77e-03 |
| 2000 | 5.74e-02 ± 0.00e+00 | 6.19e-02 ± 0.00e+00 | **7.83e-02 ± 2.23e-02** | 2.09e-02 ± 1.40e-03 | 8.09e-03 ± 3.77e-03 |

**Table S21:** Mean Top k edges F1 score (± SD) for CD34 GRNs against CD34_hematopoietic_stem_cells-derived_proerythroblasts.

| Top K edges | ctOTVelo | OTVelo | SINCERITIES | GRNBoost2 | scMTNI |
| --- | --- | --- | --- | --- | --- |
| 100 | **2.76e-02 ± 3.47e-18** | 2.67e-02 ± 0.00e+00 | 1.70e-02 ± 1.08e-02 | 2.37e-02 ± 5.98e-04 | 7.21e-03 ± 4.49e-03 |
| 200 | **5.38e-02 ± 0.00e+00** | 5.32e-02 ± 0.00e+00 | 3.43e-02 ± 2.14e-02 | 4.47e-02 ± 1.40e-03 | 9.55e-03 ± 8.86e-03 |
| 300 | **7.81e-02 ± 0.00e+00** | **7.81e-02 ± 0.00e+00** | 5.13e-02 ± 3.01e-02 | 6.55e-02 ± 1.97e-03 | 1.08e-02 ± 1.16e-02 |
| 400 | **1.02e-01 ± 1.39e-17** | 1.02e-01 ± 1.39e-17 | 6.74e-02 ± 3.94e-02 | 6.91e-02 ± 4.18e-03 | 1.13e-02 ± 1.25e-02 |
| 500 | **1.26e-01 ± 0.00e+00** | 1.25e-01 ± 0.00e+00 | 8.24e-02 ± 4.60e-02 | 6.91e-02 ± 4.18e-03 | 1.17e-02 ± 1.33e-02 |
| 1000 | 2.30e-01 ± 0.00e+00 | **2.31e-01 ± 0.00e+00** | 1.47e-01 ± 6.50e-02 | 6.91e-02 ± 4.18e-03 | 1.18e-02 ± 1.34e-02 |
| 2000 | **4.00e-01 ± 0.00e+00** | 3.96e-01 ± 0.00e+00 | 2.11e-01 ± 7.61e-02 | 6.91e-02 ± 4.18e-03 | 1.18e-02 ± 1.34e-02 |

**Table S22:**
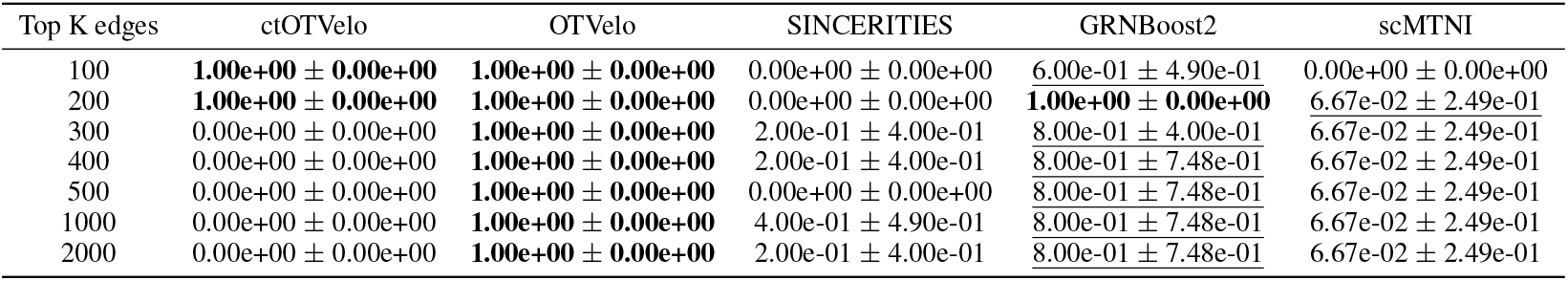
Mean Top k edges Number of predictable TFs (± SD) for Hematopoietic stem cells GRNs against CD34_hematopoietic_stem_cells_CD34CD133_hematopoietic_progenitors.

| Top K edges | ctOTVelo | OTVelo | SINCERITIES | GRNBoost2 | scMTNI |
| --- | --- | --- | --- | --- | --- |
| 100 | **1.00e+00 ± 0.00e+00** | **1.00e+00 ± 0.00e+00** | 0.00e+00 ± 0.00e+00 | 6.00e-01 ± 4.90e-01 | 0.00e+00 ± 0.00e+00 |
| 200 | **1.00e+00 ± 0.00e+00** | **1.00e+00 ± 0.00e+00** | 0.00e+00 ± 0.00e+00 | **1.00e+00 ± 0.00e+00** | 6.67e-02 ± 2.49e-01 |
| 300 | 0.00e+00 ± 0.00e+00 | **1.00e+00 ± 0.00e+00** | 2.00e-01 ± 4.00e-01 | 8.00e-01 ± 4.00e-01 | 6.67e-02 ± 2.49e-01 |
| 400 | 0.00e+00 ± 0.00e+00 | **1.00e+00 ± 0.00e+00** | 2.00e-01 ± 4.00e-01 | 8.00e-01 ± 7.48e-01 | 6.67e-02 ± 2.49e-01 |
| 500 | 0.00e+00 ± 0.00e+00 | **1.00e+00 ± 0.00e+00** | 0.00e+00 ± 0.00e+00 | 8.00e-01 ± 7.48e-01 | 6.67e-02 ± 2.49e-01 |
| 1000 | 0.00e+00 ± 0.00e+00 | **1.00e+00 ± 0.00e+00** | 4.00e-01 ± 4.90e-01 | 8.00e-01 ± 7.48e-01 | 6.67e-02 ± 2.49e-01 |
| 2000 | 0.00e+00 ± 0.00e+00 | **1.00e+00 ± 0.00e+00** | 2.00e-01 ± 4.00e-01 | 8.00e-01 ± 7.48e-01 | 6.67e-02 ± 2.49e-01 |

**Table S23:** Mean Top k edges Number of predictable TFs (± SD) for Hematopoietic stem cells GRNs against CD34_hematopoietic_stem_cells-derived_proerythroblasts.

| Top K edges | ctOTVelo | OTVelo | SINCERITIES | GRNBoost2 | scMTNI |
| --- | --- | --- | --- | --- | --- |
| 100 | 0.00e+00 ± 0.00e+00 | 0.00e+00 ± 0.00e+00 | 0.00e+00 ± 0.00e+00 | **6.00e-01 ± 4.90e-01** | 0.00e+00 ± 0.00e+00 |
| 200 | **1.00e+00 ± 0.00e+00** | **1.00e+00 ± 0.00e+00** | 0.00e+00 ± 0.00e+00 | 6.00e-01 ± 4.90e-01 | 0.00e+00 ± 0.00e+00 |
| 300 | **1.00e+00 ± 0.00e+00** | **1.00e+00 ± 0.00e+00** | 0.00e+00 ± 0.00e+00 | 4.00e-01 ± 4.90e-01 | 0.00e+00 ± 0.00e+00 |
| 400 | **1.00e+00 ± 0.00e+00** | 0.00e+00 ± 0.00e+00 | 0.00e+00 ± 0.00e+00 | 4.00e-01 ± 4.90e-01 | 0.00e+00 ± 0.00e+00 |
| 500 | **1.00e+00 ± 0.00e+00** | 0.00e+00 ± 0.00e+00 | 0.00e+00 ± 0.00e+00 | 4.00e-01 ± 4.90e-01 | 0.00e+00 ± 0.00e+00 |
| 1000 | 0.00e+00 ± 0.00e+00 | 0.00e+00 ± 0.00e+00 | 0.00e+00 ± 0.00e+00 | **4.00e-01 ± 4.90e-01** | 0.00e+00 ± 0.00e+00 |
| 2000 | 0.00e+00 ± 0.00e+00 | 0.00e+00 ± 0.00e+00 | 0.00e+00 ± 0.00e+00 | **4.00e-01 ± 4.90e-01** | 0.00e+00 ± 0.00e+00 |

**Table S24:**
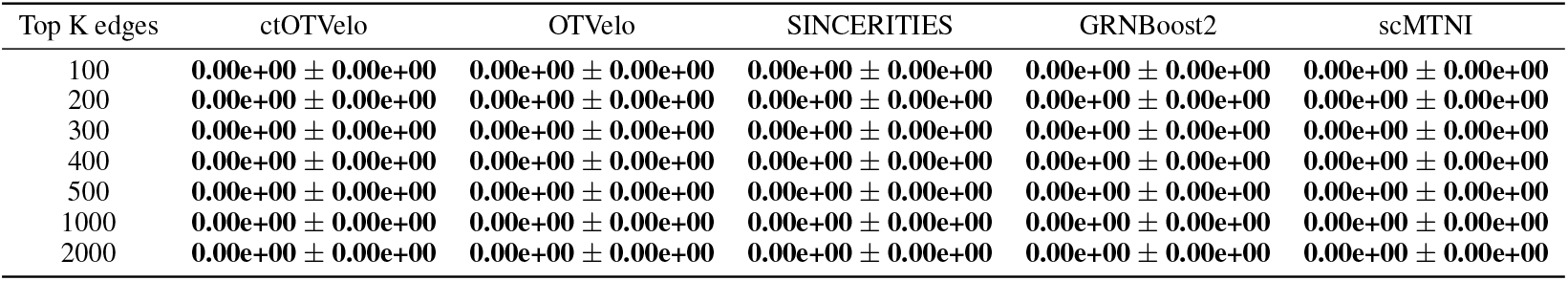
Mean Top k edges Number of predictable TFs (± SD) for Monocyte GRNs against CD14_monocytes.

**Table S25:**
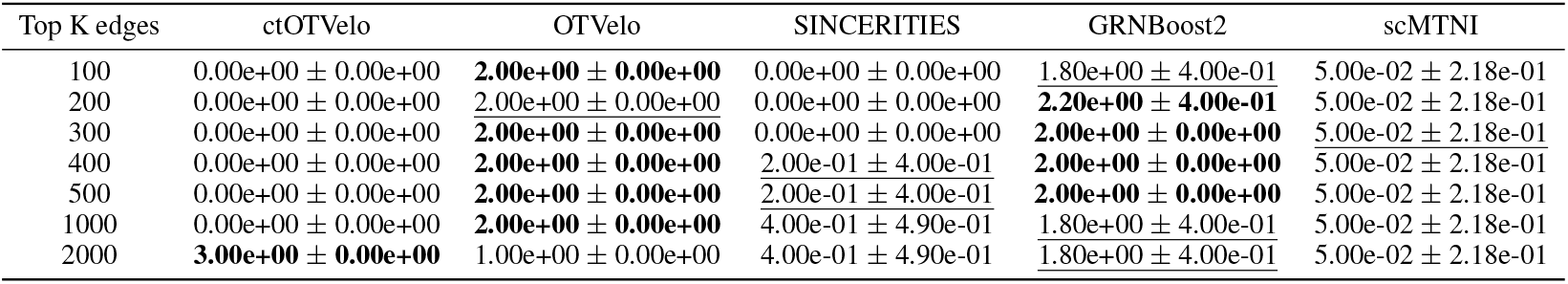
Mean Top k edges Number of predictable TFs (± SD) for Common myeloid progenitors GRNs against megakaryocytes.

**Table S26:**
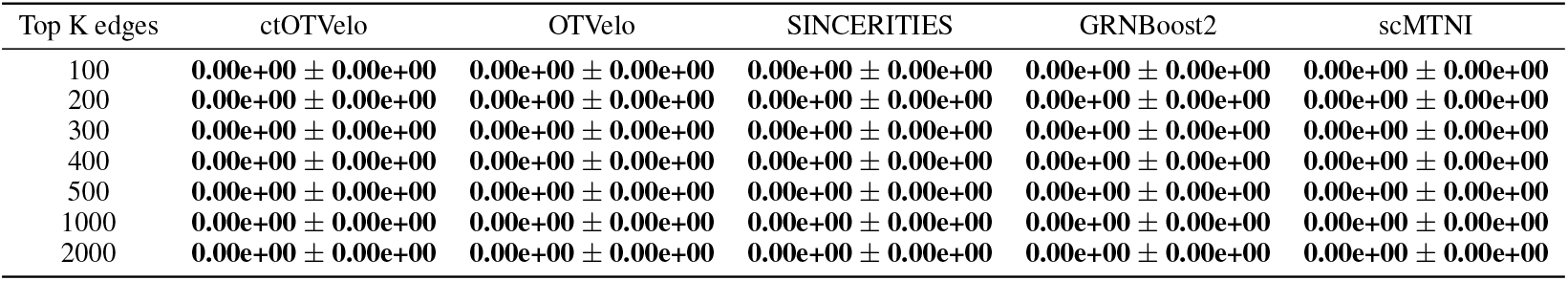
Mean Top k edges Number of predictable TFs (± SD) for Common myeloid progenitors GRNs against erythroid_progenitors.

**Table S27:**
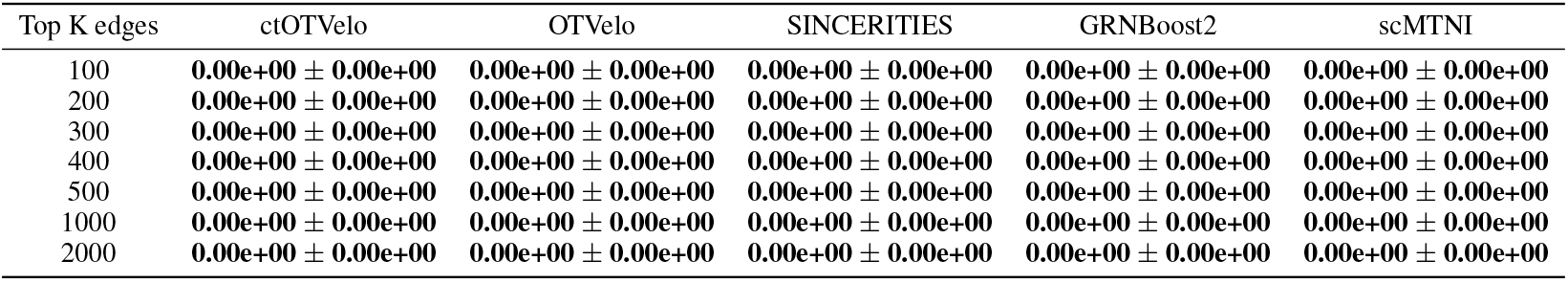
Mean Top k edges Number of predictable TFs (± SD) for Common myeloid progenitors GRNs against B-cells.

**Table S28:**
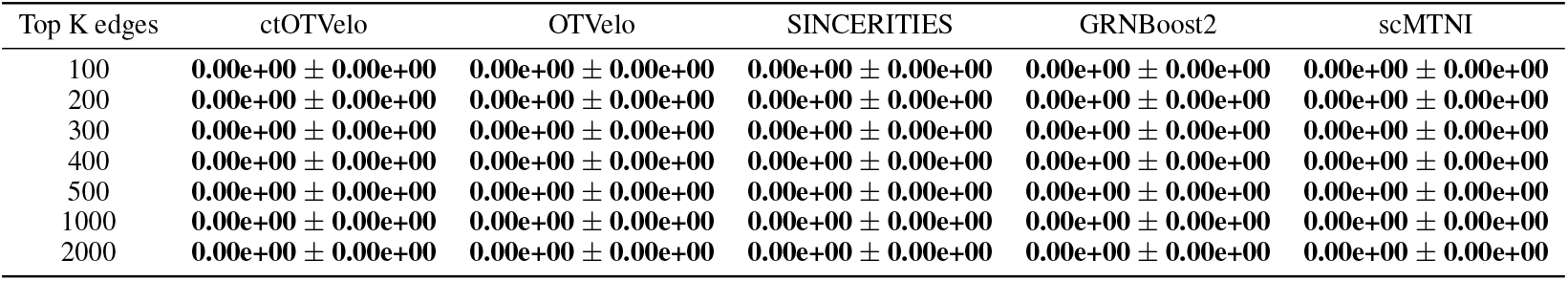
Mean Top k edges Number of predictable TFs (± SD) for Common myeloid progenitors GRNs against R3R4_erythroid_cells.

**Table S29:** Mean Top k edges Number of predictable TFs (± SD) for Granulocyte-macrophage progenitors GRNs against GM_B-cells.

| Top K edges | ctOTVelo | OTVelo | SINCERITIES | GRNBoost2 | scMTNI |
| --- | --- | --- | --- | --- | --- |
| 100 | **0.00e+00 ± 0.00e+00** | **0.00e+00 ± 0.00e+00** | **0.00e+00 ± 0.00e+00** | **0.00e+00 ± 0.00e+00** | **0.00e+00 ± 0.00e+00** |
| 200 | 0.00e+00 ± 0.00e+00 | **1.00e+00 ± 0.00e+00** | 4.00e-01 ± 4.90e-01 | 0.00e+00 ± 0.00e+00 | 0.00e+00 ± 0.00e+00 |
| 300 | 0.00e+00 ± 0.00e+00 | **1.00e+00 ± 0.00e+00** | <u>2.00e-01 ± 4.00e-01</u> | 0.00e+00 ± 0.00e+00 | 0.00e+00 ± 0.00e+00 |
| 400 | 0.00e+00 ± 0.00e+00 | **2.00e+00 ± 0.00e+00** | <u>4.00e-01 ± 4.90e-01</u> | 0.00e+00 ± 0.00e+00 | 0.00e+00 ± 0.00e+00 |
| 500 | 0.00e+00 ± 0.00e+00 | 0.00e+00 ± 0.00e+00 | **4.00e-01 ± 4.90e-01** | 0.00e+00 ± 0.00e+00 | 0.00e+00 ± 0.00e+00 |
| 1000 | 0.00e+00 ± 0.00e+00 | **2.00e+00 ± 0.00e+00** | 8.00e-01 ± 1.17e+00 | 0.00e+00 ± 0.00e+00 | 0.00e+00 ± 0.00e+00 |
| 2000 | 0.00e+00 ± 0.00e+00 | <u>1.00e+00 ± 0.00e+00</u> | **1.40e+00 ± 1.02e+00** | 4.00e-01 ± 8.00e-01 | 0.00e+00 ± 0.00e+00 |

**Table S30:** Mean Top k edges Number of predictable TFs (± SD) for CD34 GRNs against CD34_hematopoietic_stem_cells_CD34CD133_hematopoietic_progenitors.

| Top K edges | ctOTVelo | OTVelo | SINCERITIES | GRNBoost2 | scMTNI |
| --- | --- | --- | --- | --- | --- |
| 100 | **1.00e+00 ± 0.00e+00** | **1.00e+00 ± 0.00e+00** | 0.00e+00 ± 0.00e+00 | 6.00e-01 ± 4.90e-01 | 0.00e+00 ± 0.00e+00 |
| 200 | **1.00e+00 ± 0.00e+00** | **1.00e+00 ± 0.00e+00** | 0.00e+00 ± 0.00e+00 | **1.00e+00 ± 0.00e+00** | <u>6.67e-02 ± 2.49e-01</u> |
| 300 | **1.00e+00 ± 0.00e+00** | **1.00e+00 ± 0.00e+00** | 2.00e-01 ± 4.00e-01 | <u>8.00e-01 ± 4.00e-01</u> | 6.67e-02 ± 2.49e-01 |
| 400 | **1.00e+00 ± 0.00e+00** | **1.00e+00 ± 0.00e+00** | 2.00e-01 ± 4.00e-01 | <u>8.00e-01 ± 7.48e-01</u> | 6.67e-02 ± 2.49e-01 |
| 500 | **1.00e+00 ± 0.00e+00** | **1.00e+00 ± 0.00e+00** | 0.00e+00 ± 0.00e+00 | <u>8.00e-01 ± 7.48e-01</u> | 6.67e-02 ± 2.49e-01 |
| 1000 | **1.00e+00 ± 0.00e+00** | **1.00e+00 ± 0.00e+00** | 4.00e-01 ± 4.90e-01 | <u>8.00e-01 ± 7.48e-01</u> | 6.67e-02 ± 2.49e-01 |
| 2000 | **1.00e+00 ± 0.00e+00** | **1.00e+00 ± 0.00e+00** | 2.00e-01 ± 4.00e-01 | <u>8.00e-01 ± 7.48e-01</u> | 6.67e-02 ± 2.49e-01 |

**Table S31:**
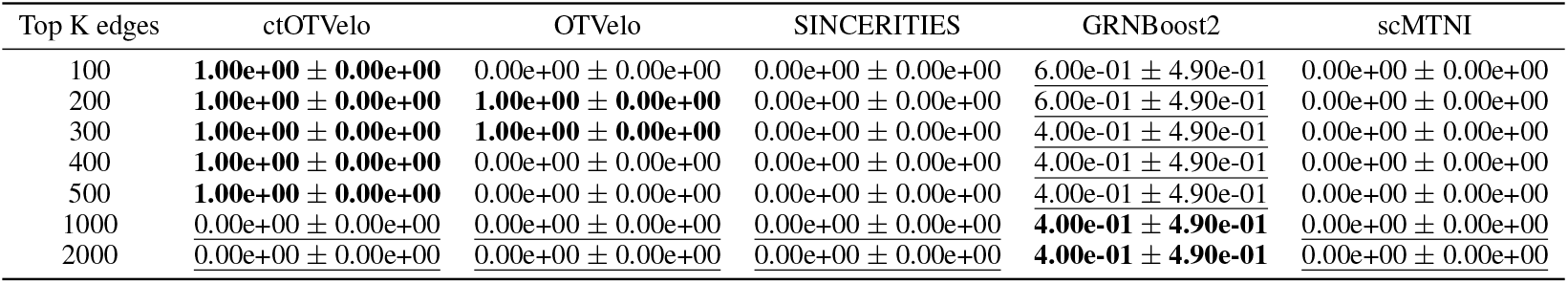
Mean Top k edges Number of predictable TFs (± SD) for CD34 GRNs against CD34_hematopoietic_stem_cells-derived_proerythroblasts.

| Top K edges | ctOTVelo | OTVelo | SINCERITIES | GRNBoost2 | scMTNI |
| --- | --- | --- | --- | --- | --- |
| 100 | **1.00e+00 ± 0.00e+00** | 0.00e+00 ± 0.00e+00 | 0.00e+00 ± 0.00e+00 | <u>6.00e-01 ± 4.90e-01</u> | 0.00e+00 ± 0.00e+00 |
| 200 | **1.00e+00 ± 0.00e+00** | **1.00e+00 ± 0.00e+00** | 0.00e+00 ± 0.00e+00 | <u>6.00e-01 ± 4.90e-01</u> | 0.00e+00 ± 0.00e+00 |
| 300 | **1.00e+00 ± 0.00e+00** | **1.00e+00 ± 0.00e+00** | 0.00e+00 ± 0.00e+00 | <u>4.00e-01 ± 4.90e-01</u> | 0.00e+00 ± 0.00e+00 |
| 400 | **1.00e+00 ± 0.00e+00** | 0.00e+00 ± 0.00e+00 | 0.00e+00 ± 0.00e+00 | <u>4.00e-01 ± 4.90e-01</u> | 0.00e+00 ± 0.00e+00 |
| 500 | **1.00e+00 ± 0.00e+00** | 0.00e+00 ± 0.00e+00 | 0.00e+00 ± 0.00e+00 | <u>4.00e-01 ± 4.90e-01</u> | 0.00e+00 ± 0.00e+00 |
| 1000 | 0.00e+00 ± 0.00e+00 | 0.00e+00 ± 0.00e+00 | 0.00e+00 ± 0.00e+00 | **4.00e-01 ± 4.90e-01** | 0.00e+00 ± 0.00e+00 |
| 2000 | <u>0.00e+00 ± 0.00e+00</u> | <u>0.00e+00 ± 0.00e+00</u> | <u>0.00e+00 ± 0.00e+00</u> | **4.00e-01 ± 4.90e-01** | <u>0.00e+00 ± 0.00e+00</u> |

## C Full quantitative metrics for mouse organogenesis

**Table S32:**
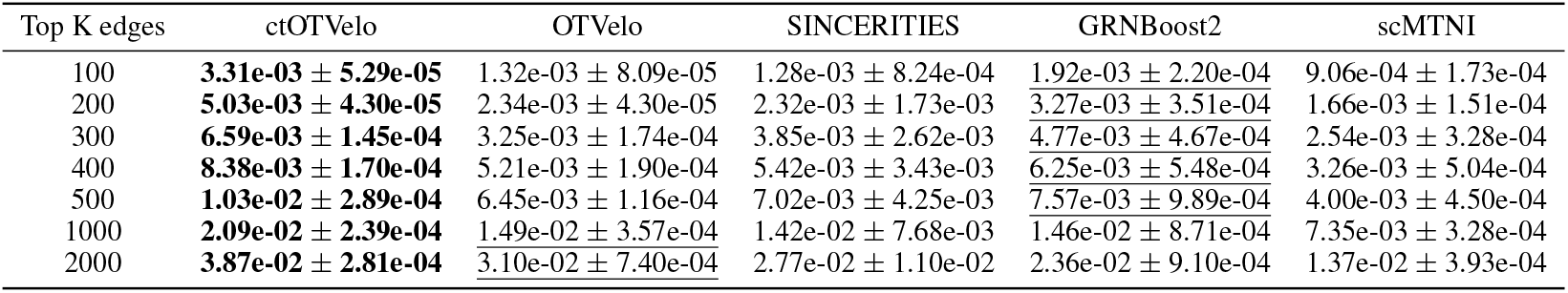
Mean Top k edges F1 score (± SD) for global GRNs against mESC_chipunion.

**Table S33:**
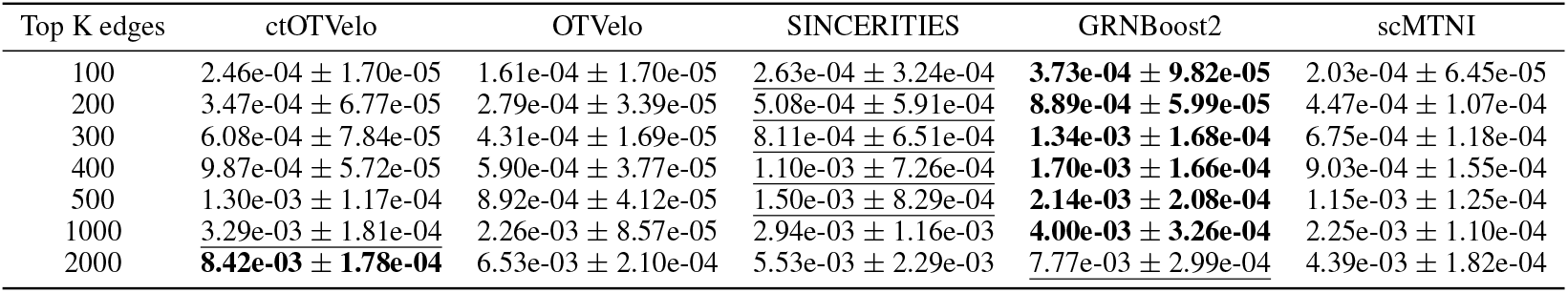
Mean Top k edges F1 score (± SD) for global GRNs against mESC_KDUnion.

**Table S34:** Mean Top k edges F1 score (± SD) for global GRNs against mESC_chipunion_KDUnion_intersect.

| Top K edges | ctOTVelo | OTVelo | SINCERITIES | GRNBoost2 | scMTNI |
| --- | --- | --- | --- | --- | --- |
| 100 | **1.94e-02 ± 5.11e-04** | 5.49e-03 ± 3.23e-04 | 6.78e-03 ± 5.27e-03 | <u>1.42e-02 ± 2.08e-03</u> | 7.20e-03 ± 1.48e-03 |
| 200 | **3.65e-02 ± 6.94e-04** | 1.65e-02 ± 3.10e-04 | 1.16e-02 ± 6.82e-03 | <u>2.30e-02 ± 2.43e-03</u> | 1.26e-02 ± 2.55e-03 |
| 300 | **5.11e-02 ± 7.62e-04** | 2.81e-02 ± 1.21e-03 | 1.34e-02 ± 8.67e-03 | <u>2.86e-02 ± 3.14e-03</u> | 1.51e-02 ± 2.00e-03 |
| 400 | **6.28e-02 ± 1.84e-03** | <u>3.66e-02 ± 1.40e-03</u> | 1.67e-02 ± 7.45e-03 | <u>2.88e-02 ± 3.37e-03</u> | 1.51e-02 ± 2.00e-03 |
| 500 | **7.55e-02 ± 3.28e-03** | <u>4.48e-02 ± 8.34e-04</u> | 1.88e-02 ± 6.25e-03 | 2.88e-02 ± 3.37e-03 | 1.51e-02 ± 2.00e-03 |
| 1000 | **1.30e-01 ± 1.73e-03** | <u>9.70e-02 ± 2.52e-03</u> | 2.98e-02 ± 5.34e-03 | 2.88e-02 ± 3.37e-03 | 1.51e-02 ± 2.00e-03 |
| 2000 | **1.70e-01 ± 1.60e-03** | <u>1.46e-01 ± 2.80e-03</u> | 4.80e-02 ± 8.06e-03 | 2.88e-02 ± 3.37e-03 | 1.51e-02 ± 2.00e-03 |

**Table S35:** Mean Top k edges Precision (± SD) for global GRNs against mESC_chipunion.

| Top K edges | ctOTVelo | OTVelo | SINCERITIES | GRNBoost2 | scMTNI |
| --- | --- | --- | --- | --- | --- |
| 100 | **3.06e-01 ± 4.90e-03** | 1.22e-01 ± 7.48e-03 | 1.18e-01 ± 7.63e-02 | 1.78e-01 ± 2.04e-02 | <u>1.90e-01 ± 3.63e-02</u> |
| 200 | **2.34e-01 ± 2.00e-03** | 1.09e-01 ± 2.00e-03 | 1.08e-01 ± 8.05e-02 | 1.52e-01 ± 1.63e-02 | <u>1.74e-01 ± 1.59e-02</u> |
| 300 | **2.05e-01 ± 4.52e-03** | 1.01e-01 ± 5.42e-03 | 1.20e-01 ± 8.17e-02 | 1.49e-01 ± 1.45e-02 | <u>1.78e-01 ± 2.31e-02</u> |
| 400 | **1.97e-01 ± 4.00e-03** | 1.23e-01 ± 4.47e-03 | 1.28e-01 ± 8.06e-02 | 1.47e-01 ± 1.29e-02 | <u>1.72e-01 ± 2.67e-02</u> |
| 500 | **1.95e-01 ± 5.46e-03** | 1.22e-01 ± 2.19e-03 | 1.33e-01 ± 8.03e-02 | 1.43e-01 ± 1.87e-02 | <u>1.69e-01 ± 1.91e-02</u> |
| 1000 | **2.03e-01 ± 2.32e-03** | 1.45e-01 ± 3.46e-03 | 1.38e-01 ± 7.46e-02 | 1.42e-01 ± 8.45e-03 | <u>1.57e-01 ± 7.03e-03</u> |
| 2000 | **1.97e-01 ± 1.44e-03** | <u>1.58e-01 ± 3.77e-03</u> | 1.41e-01 ± 5.61e-02 | 1.39e-01 ± 5.39e-03 | 1.50e-01 ± 4.36e-03 |

**Table S36:** Mean Top k edges Precision (± SD) for global GRNs against mESC_KDUnion.

| Top K edges | ctOTVelo | OTVelo | SINCERITIES | GRNBoost2 | scMTNI |
| --- | --- | --- | --- | --- | --- |
| 100 | 5.80e-02 ± 4.00e-03 | 3.80e-02 ± 4.00e-03 | 6.20e-02 ± 7.63e-02 | **8.80e-02 ± 2.32e-02** | <u>7.60e-02 ± 2.42e-02</u> |
| 200 | 4.10e-02 ± 8.00e-03 | 3.30e-02 ± 4.00e-03 | 6.00e-02 ± 6.98e-02 | **1.05e-01 ± 7.07e-03** | <u>8.40e-02 ± 2.01e-02</u> |
| 300 | 4.80e-02 ± 6.18e-03 | 3.40e-02 ± 1.33e-03 | 6.40e-02 ± 5.14e-02 | **1.06e-01 ± 1.32e-02** | <u>8.47e-02 ± 1.48e-02</u> |
| 400 | 5.85e-02 ± 3.39e-03 | 3.50e-02 ± 2.24e-03 | 6.55e-02 ± 4.31e-02 | **1.01e-01 ± 9.82e-03** | <u>8.50e-02 ± 1.47e-02</u> |
| 500 | 6.16e-02 ± 5.57e-03 | 4.24e-02 ± 1.96e-03 | 7.12e-02 ± 3.94e-02 | **1.02e-01 ± 9.91e-03** | <u>8.68e-02 ± 9.68e-03</u> |
| 1000 | 7.90e-02 ± 4.34e-03 | 5.44e-02 ± 2.06e-03 | 7.06e-02 ± 2.80e-02 | **9.62e-02 ± 7.83e-03** | <u>8.54e-02 ± 4.27e-03</u> |
| 2000 | **1.03e-01 ± 2.18e-03** | 8.00e-02 ± 2.57e-03 | 6.78e-02 ± 2.81e-02 | <u>9.52e-02 ± 3.67e-03</u> | 8.45e-02 ± 3.73e-03 |

**Table S37:** Mean Top k edges Precision (± SD) for global GRNs against mESC_chipunion_KDUnion_intersect.

| Top K edges | ctOTVelo | OTVelo | SINCERITIES | GRNBoost2 | scMTNI |
| --- | --- | --- | --- | --- | --- |
| 100 | **2.40e-01 ± 6.32e-03** | 6.80e-02 ± 4.00e-03 | 8.40e-02 ± 6.53e-02 | <u>1.76e-01 ± 2.58e-02</u> | 1.46e-01 ± 3.01e-02 |
| 200 | **2.35e-01 ± 4.47e-03** | 1.06e-01 ± 2.00e-03 | 7.50e-02 ± 4.39e-02 | <u>1.48e-01 ± 1.57e-02</u> | 1.31e-01 ± 2.65e-02 |
| 300 | **2.28e-01 ± 3.40e-03** | 1.25e-01 ± 5.42e-03 | 6.00e-02 ± 3.87e-02 | <u>1.29e-01 ± 1.27e-02</u> | 1.21e-01 ± 2.20e-02 |
| 400 | **2.18e-01 ± 6.40e-03** | <u>1.27e-01 ± 4.85e-03</u> | 5.80e-02 ± 2.59e-02 | 1.19e-01 ± 1.01e-02 | 1.20e-01 ± 2.29e-02 |
| 500 | **2.17e-01 ± 9.43e-03** | <u>1.29e-01 ± 2.40e-03</u> | 5.40e-02 ± 1.80e-02 | 1.19e-01 ± 1.01e-02 | 1.20e-01 ± 2.29e-02 |
| 1000 | **2.20e-01 ± 2.93e-03** | <u>1.64e-01 ± 4.26e-03</u> | 5.04e-02 ± 9.02e-03 | 1.19e-01 ± 1.01e-02 | 1.20e-01 ± 2.29e-02 |
| 2000 | **1.86e-01 ± 1.75e-03** | <u>1.59e-01 ± 3.06e-03</u> | 5.25e-02 ± 8.81e-03 | 1.19e-01 ± 1.01e-02 | 1.20e-01 ± 2.29e-02 |

**Table S38:** Mean Top k edges Recall (± SD) for global GRNs against mESC_chipunion.

| Top K edges | ctOTVelo | OTVelo | SINCERITIES | GRNBoost2 | scMTNI |
| --- | --- | --- | --- | --- | --- |
| 100 | **1.66e-03 ± 2.66e-05** | 6.63e-04 ± 4.07e-05 | 6.41e-04 ± 4.14e-04 | <u>9.67e-04 ± 1.11e-04</u> | 4.54e-04 ± 8.67e-05 |
| 200 | **2.54e-03 ± 2.17e-05** | 1.18e-03 ± 2.17e-05 | 1.17e-03 ± 8.74e-04 | <u>1.65e-03 ± 1.77e-04</u> | 8.32e-04 ± 7.58e-05 |
| 300 | **3.35e-03 ± 7.37e-05** | 1.65e-03 ± 8.83e-05 | 1.96e-03 ± 1.33e-03 | <u>2.42e-03 ± 2.37e-04</u> | 1.28e-03 ± 1.65e-04 |
| 400 | **4.28e-03 ± 8.69e-05** | 2.66e-03 ± 9.72e-05 | 2.77e-03 ± 1.75e-03 | <u>3.19e-03 ± 2.80e-04</u> | 1.64e-03 ± 2.55e-04 |
| 500 | **5.30e-03 ± 1.48e-04** | 3.31e-03 ± 5.95e-05 | 3.61e-03 ± 2.18e-03 | <u>3.89e-03 ± 5.08e-04</u> | 2.02e-03 ± 2.28e-04 |
| 1000 | **1.10e-02 ± 1.26e-04** | 7.88e-03 ± 1.88e-04 | 7.51e-03 ± 4.05e-03 | 7.70e-03 ± 4.59e-04 | 3.76e-03 ± 1.68e-04 |
| 2000 | **2.14e-02 ± 1.56e-04** | <u>1.72e-02 ± 4.10e-04</u> | 1.53e-02 ± 6.09e-03 | 1.29e-02 ± 5.06e-04 | 7.18e-03 ± 2.06e-04 |

**Table S39:** Mean Top k edges Recall (± SD) for global GRNs against mESC_KDUnion.

| Top K edges | ctOTVelo | OTVelo | SINCERITIES | GRNBoost2 | scMTNI |
| --- | --- | --- | --- | --- | --- |
| 100 | 1.23e-04 ± 8.50e-06 | 8.08e-05 ± 8.50e-06 | 1.32e-04 ± 1.62e-04 | **1.87e-04 ± 4.92e-05** | 1.01e-04 ± 3.23e-05 |
| 200 | 1.74e-04 ± 3.40e-05 | 1.40e-04 ± 1.70e-05 | 2.55e-04 ± 2.97e-04 | **4.46e-04 ± 3.01e-05** | 2.24e-04 ± 5.36e-05 |
| 300 | 3.06e-04 ± 3.94e-05 | 2.17e-04 ± 8.50e-06 | 4.08e-04 ± 3.28e-04 | **6.76e-04 ± 8.44e-05** | 3.39e-04 ± 5.92e-05 |
| 400 | 4.97e-04 ± 2.88e-05 | 2.98e-04 ± 1.90e-05 | 5.57e-04 ± 3.66e-04 | **8.59e-04 ± 8.35e-05** | 4.54e-04 ± 7.79e-05 |
| 500 | 6.55e-04 ± 5.92e-05 | 4.51e-04 ± 2.08e-05 | 7.57e-04 ± 4.19e-04 | **1.08e-03 ± 1.05e-04** | 5.79e-04 ± 6.30e-05 |
| 1000 | 1.68e-03 ± 9.22e-05 | 1.16e-03 ± 4.38e-05 | 1.50e-03 ± 5.95e-04 | **2.05e-03 ± 1.67e-04** | 1.14e-03 ± 5.57e-05 |
| 2000 | **4.39e-03 ± 9.28e-05** | 3.40e-03 ± 1.09e-04 | 2.88e-03 ± 1.19e-03 | 4.05e-03 ± 1.56e-04 | 2.26e-03 ± 9.30e-05 |

**Table S40:** Mean Top k edges Recall (± SD) for global GRNs against mESC_chipunion_KDUnion_intersect.

| Top K edges | ctOTVelo | OTVelo | SINCERITIES | GRNBoost2 | scMTNI |
| --- | --- | --- | --- | --- | --- |
| 100 | **1.01e-02 ± 2.66e-04** | 2.86e-03 ± 1.68e-04 | 3.53e-03 ± 2.75e-03 | 7.40e-03 ± 1.08e-03 | 3.69e-03 ± 7.60e-04 |
| 200 | **1.98e-02 ± 3.76e-04** | 8.92e-03 ± 1.68e-04 | 6.31e-03 ± 3.70e-03 | 1.25e-02 ± 1.32e-03 | 6.62e-03 ± 1.34e-03 |
| 300 | **2.88e-02 ± 4.29e-04** | 1.58e-02 ± 6.84e-04 | 7.57e-03 ± 4.88e-03 | 1.61e-02 ± 1.79e-03 | 8.04e-03 ± 1.05e-03 |
| 400 | **3.67e-02 ± 1.08e-03** | 2.14e-02 ± 8.16e-04 | 9.76e-03 ± 4.35e-03 | 1.64e-02 ± 2.01e-03 | 8.04e-03 ± 1.05e-03 |
| 500 | **4.57e-02 ± 1.98e-03** | 2.71e-02 ± 5.05e-04 | 1.14e-02 ± 3.78e-03 | 1.64e-02 ± 2.01e-03 | 8.04e-03 ± 1.05e-03 |
| 1000 | **9.26e-02 ± 1.23e-03** | 6.89e-02 ± 1.79e-03 | 2.12e-02 ± 3.80e-03 | 1.64e-02 ± 2.01e-03 | 8.04e-03 ± 1.05e-03 |
| 2000 | **1.56e-01 ± 1.47e-03** | 1.34e-01 ± 2.57e-03 | 4.42e-02 ± 7.42e-03 | 1.64e-02 ± 2.01e-03 | 8.04e-03 ± 1.05e-03 |

**Table S41:** Mean Top k edges Number of predictable TFs (± SD) for global GRNs against mESC_chipunion.

| Top K edges | ctOTVelo | OTVelo | SINCERITIES | GRNBoost2 | scMTNI |
| --- | --- | --- | --- | --- | --- |
| 100 | **1.00e+00 ± 0.00e+00** | **1.00e+00 ± 1.73e+00** | 2.00e-01 ± 4.00e-01 | 0.00e+00 ± 0.00e+00 | 2.00e-01 ± 4.00e-01 |
| 200 | **1.00e+00 ± 0.00e+00** | 4.00e-01 ± 7.35e-01 | 6.00e-01 ± 8.00e-01 | 0.00e+00 ± 0.00e+00 | 2.00e-01 ± 4.00e-01 |
| 300 | **1.00e+00 ± 0.00e+00** | 6.50e-01 ± 1.15e+00 | 6.00e-01 ± 8.00e-01 | 0.00e+00 ± 0.00e+00 | 2.00e-01 ± 4.00e-01 |
| 400 | 8.00e-01 ± 4.00e-01 | 7.50e-01 ± 1.30e+00 | **1.20e+00 ± 7.48e-01** | 0.00e+00 ± 0.00e+00 | 2.00e-01 ± 4.00e-01 |
| 500 | 2.00e-01 ± 4.00e-01 | 8.00e-01 ± 1.40e+00 | **1.20e+00 ± 7.48e-01** | 0.00e+00 ± 0.00e+00 | 2.00e-01 ± 4.00e-01 |
| 1000 | 1.00e+00 ± 0.00e+00 | 1.05e+00 ± 1.83e+00 | **1.20e+00 ± 9.80e-01** | 0.00e+00 ± 0.00e+00 | 0.00e+00 ± 0.00e+00 |
| 2000 | 1.00e+00 ± 0.00e+00 | 1.40e+00 ± 2.01e+00 | **2.20e+00 ± 1.17e+00** | 2.00e-01 ± 4.00e-01 | 0.00e+00 ± 0.00e+00 |

**Table S42:** Mean Top k edges Number of predictable TFs (± SD) for global GRNs against mESC_KDUnion.

| Top K edges | ctOTVelo | OTVelo | SINCERITIES | GRNBoost2 | scMTNI |
| --- | --- | --- | --- | --- | --- |
| 100 | 0.00e+00 ± 0.00e+00 | **1.00e+00 ± 1.73e+00** | 0.00e+00 ± 0.00e+00 | 0.00e+00 ± 0.00e+00 | 0.00e+00 ± 0.00e+00 |
| 200 | 0.00e+00 ± 0.00e+00 | **4.00e-01 ± 7.35e-01** | **4.00e-01 ± 4.90e-01** | 0.00e+00 ± 0.00e+00 | 0.00e+00 ± 0.00e+00 |
| 300 | 0.00e+00 ± 0.00e+00 | **6.50e-01 ± 1.15e+00** | 2.00e-01 ± 4.00e-01 | 0.00e+00 ± 0.00e+00 | 0.00e+00 ± 0.00e+00 |
| 400 | 0.00e+00 ± 0.00e+00 | **7.50e-01 ± 1.30e+00** | 2.00e-01 ± 4.00e-01 | 2.00e-01 ± 4.00e-01 | 0.00e+00 ± 0.00e+00 |
| 500 | 0.00e+00 ± 0.00e+00 | **8.00e-01 ± 1.40e+00** | 4.00e-01 ± 4.90e-01 | 2.00e-01 ± 4.00e-01 | 0.00e+00 ± 0.00e+00 |
| 1000 | 0.00e+00 ± 0.00e+00 | **1.05e+00 ± 1.83e+00** | 4.00e-01 ± 4.90e-01 | 2.00e-01 ± 4.00e-01 | 0.00e+00 ± 0.00e+00 |
| 2000 | 8.00e-01 ± 4.00e-01 | **1.40e+00 ± 2.01e+00** | 4.00e-01 ± 4.90e-01 | 2.00e-01 ± 4.00e-01 | 0.00e+00 ± 0.00e+00 |

**Table S43:**
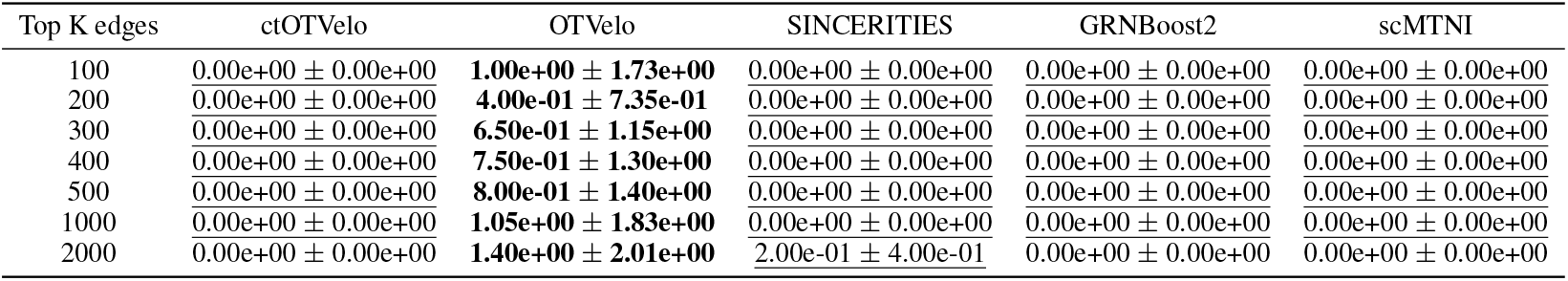
Mean Top k edges Number of predictable TFs (± SD) for global GRNs against mESC_chipunion_KDUnion_intersect.

